# SARS-CoV-2 Omicron efficiently infects human airway, but not alveolar epithelium

**DOI:** 10.1101/2022.01.19.476898

**Authors:** Mart M. Lamers, Anna Z. Mykytyn, Tim I. Breugem, Nathalie Groen, Kèvin Knoops, Debby Schipper, Romy van Acker, Petra B. van den Doel, Theo Bestebroer, Charlotte D. Koopman, Chantal Reusken, Mauro J. Muraro, Corine H. GeurtsvanKessel, Martin E. van Royen, Peter J. Peters, Jingshu Zhang, Bart L. Haagmans

**Author notes:** These authors contributed equally (Mart M. Lamers, Anna Z. Mykytyn, Tim I. Breugem).

## Abstract

In late 2021, the highly mutated SARS-CoV-2 Omicron variant emerged, raising concerns about its potential extensive immune evasion, increased transmissibility and pathogenicity. Here, we used organoids of the human airways and alveoli to investigate Omicron’s fitness and replicative potential in comparison with earlier SARS-CoV-2 variants. We report that Omicron replicates more rapidly in the airways and has an increased fitness compared to the early 614G variant and Delta. In contrast, Omicron did not replicate productively in human alveolar type 2 cells. Mechanistically, we show that Omicron does not efficiently use TMPRSS2 for entry or spread through cell-cell fusion. Altogether, our data show that Omicron has an altered tropism and protease usage, potentially explaining its higher transmissibility and decreased pathogenicity.

## Main text

The devastating COVID-19 pandemic has been responsible for over 5 million deaths and continues to cause an immense global health crisis. The causative agent, SARS-CoV-2, emerged in Wuhan, China, in late 2019. Although SARS-CoV-2 was genetically and antigenically relatively stable during the first months of circulation in humans, variants that were more transmissible and/or escaped immunity emerged in late 2020 (*1*). The first variant of concern (VOC) was the Alpha variant (clade B.1.1.7) which became the dominant variant globally in early 2021. Several months later, the Delta variant (clade B.1.617.2) emerged, which had outcompeted Alpha in most countries in the summer of 2021. Epidemiological data suggest that Alpha spread 35-100% faster than its ancestral lineage and was associated with higher viral loads (*2-5*). In turn, Delta was estimated to be 55% more transmissible compared with Alpha (*6, 7*) and shown to partially escape neutralization *in vitro (8)*. Omicron (clade B.1.1.529, BA-1 and BA-2) was first detected in November 2021 in South Africa, Botswana and in a person traveling from South Africa to Hong Kong (*9-11*). Shortly thereafter, the variant was detected in several European, African and Asian countries as well as the USA (*9, 10, 12, 13*). Currently, Omicron appears to be rapidly replacing Delta in many countries, indicating it may transmit more efficiently (*10, 14-16*). The spike (S) protein of Omicron harbors over 30 mutations (*10*), whereas the other VOCs only contained 9 to 12. Many of these mutations are in sites known to be involved in antigenic drift, raising concerns for extensive immune evasion. Indeed, recent studies show that Omicron efficiently evades antibody neutralization, likely compromising protection by antibodies induced upon infection or vaccination (*11, 17-25*). However, little is currently known about Omicron’s transmissibility and pathogenicity. Here, we used human airway and alveolar organoids to characterize the shedding, fitness, and virulence of SARS-CoV-2 VOCs. We report that Omicron efficiently replicates in the human airway, but not in the alveoli, and that it does not efficiently use TMPRSS2 for entry.

### Omicron efficiently replicates in airway cells, and VOCs exhibit prolonged shedding kinetics compared with the ancestral 614G virus

To faithfully recapitulate SARS-CoV-2 infection of the human airways, we used a 2D organoid-based air-liquid interface airway model described previously (*26-28*) (**Fig. 1A**). As humans shed infectious SARS-CoV-2 for approximately 8 days (*29*), we assessed the shedding kinetics of a 614D and 614G ancestral virus, Alpha, Delta and Omicron for 10 days. All viruses were propagated on Calu-3 cells to prevent culture adaptation (*27*). Virus replication was quantified by RT-qPCR and by infectious virus titration on Calu-3 cells using a plaque assay. Despite a 3.5 and 4-fold lower infectivity of Omicron compared with 614G and Delta, respectively, the infectivity on Calu-3 cells was the least variable between viruses compared with VeroE6 and VeroE6-TMPRSS2 cells (**Fig. S1A-B**) on which Omicron was severely attenuated (**Fig. S1C-F**). In the airway organoid model, we observed that the ancestral 614G virus reached its peak titer at around day 2-4 (∼10^8^ RNA copies/ml; ∼10^6^PFU/ml), after which shedding decreased by ∼100 fold between day 4 and 10 (**Fig. 1B-C**). Whereas the Alpha variant reached similar peak titers, it continued to shed high levels of infectious virus up to day 10 post-infection. Very similar shedding kinetics were observed for Delta (**Fig. 1D-E**), suggesting that both variants have evolved to shed high levels of virus for prolonged time.

**Fig. 1.**
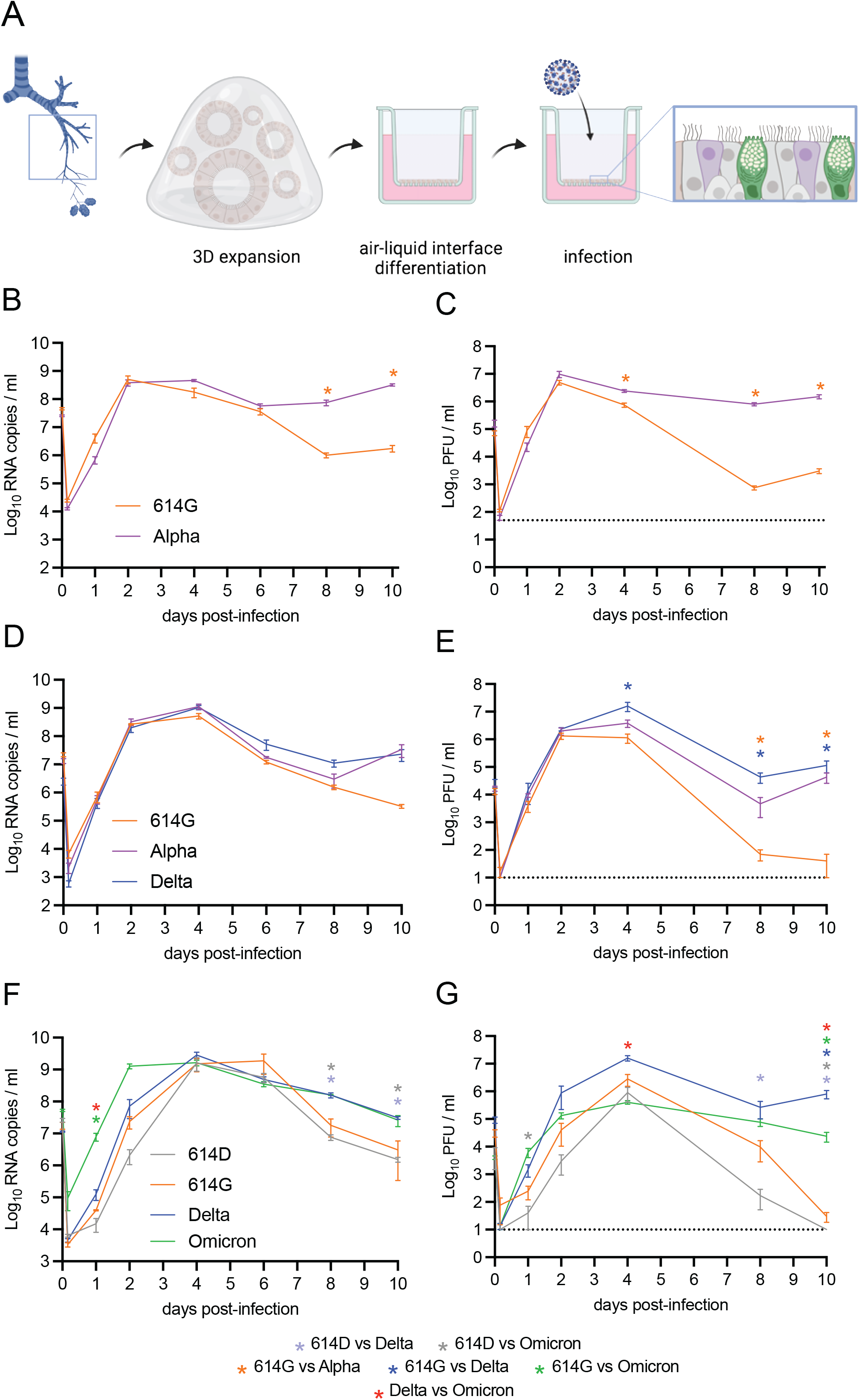
Omicron replicates rapidly in air-liquid interface airway organoids. (a) Graphical depiction of the isolation, 3D propagation and 2D air-liquid interface differentiation of human airway organoids used for SARS-CoV-2 infections. (b-c) Replication kinetics in terms of RNA (b) and PFU (c) of an ancestral 614G virus and Alpha on 2D human airway organoids. (d-e) Replication kinetics in terms of RNA (d) and PFU (e) of an ancestral 614G virus, Alpha and Delta on 2D human airway organoids. (f-g) Replication kinetics in terms of RNA (f) and PFU (g) of an ancestral 614D and 614G virus, Delta and Omicron on 2D human airway organoids. PFU = plaque forming units. Groups were compared by two-way ANOVA. *p < 0.05. Error bars depict SEM. Panel (a) was created with BioRender.com.

We subsequently characterized the shedding kinetics of Omicron in comparison to 614D, 614G, and Delta. We confirmed that Delta shed extendedly, whereas the earlier 614D and 614G viruses did not (**Fig. 1F-G**). In the same experiment, Omicron also exhibited prolonged shedding, suggesting that this is a common adaptation for all VOCs. In addition, Omicron replicated to higher titers in the 1 to 2 first days compared with Delta. This difference was best observed by looking at the RNA copies titer, as the lower infectivity of Omicron on Calu-3 cells likely caused Omicron’s live virus titers to be underestimated.

By confocal microscopy we observed that Omicron and Delta infected predominantly ciliated cells, similar to previous reports on 614G (**Fig. 2A**) (*26-28, 30, 31*). We confirmed that at day 2 post-infection, Omicron infected more cells than Delta (**Fig. 2B, D**), whereas higher infection rates were observed at day 3 for Delta (**Fig. 2C, E**). At day 10 post-infection, similar levels of infection were seen for Delta and Omicron, whereas 614D and 614G were both cleared (**Fig. 2F**).

**Fig. 2.**
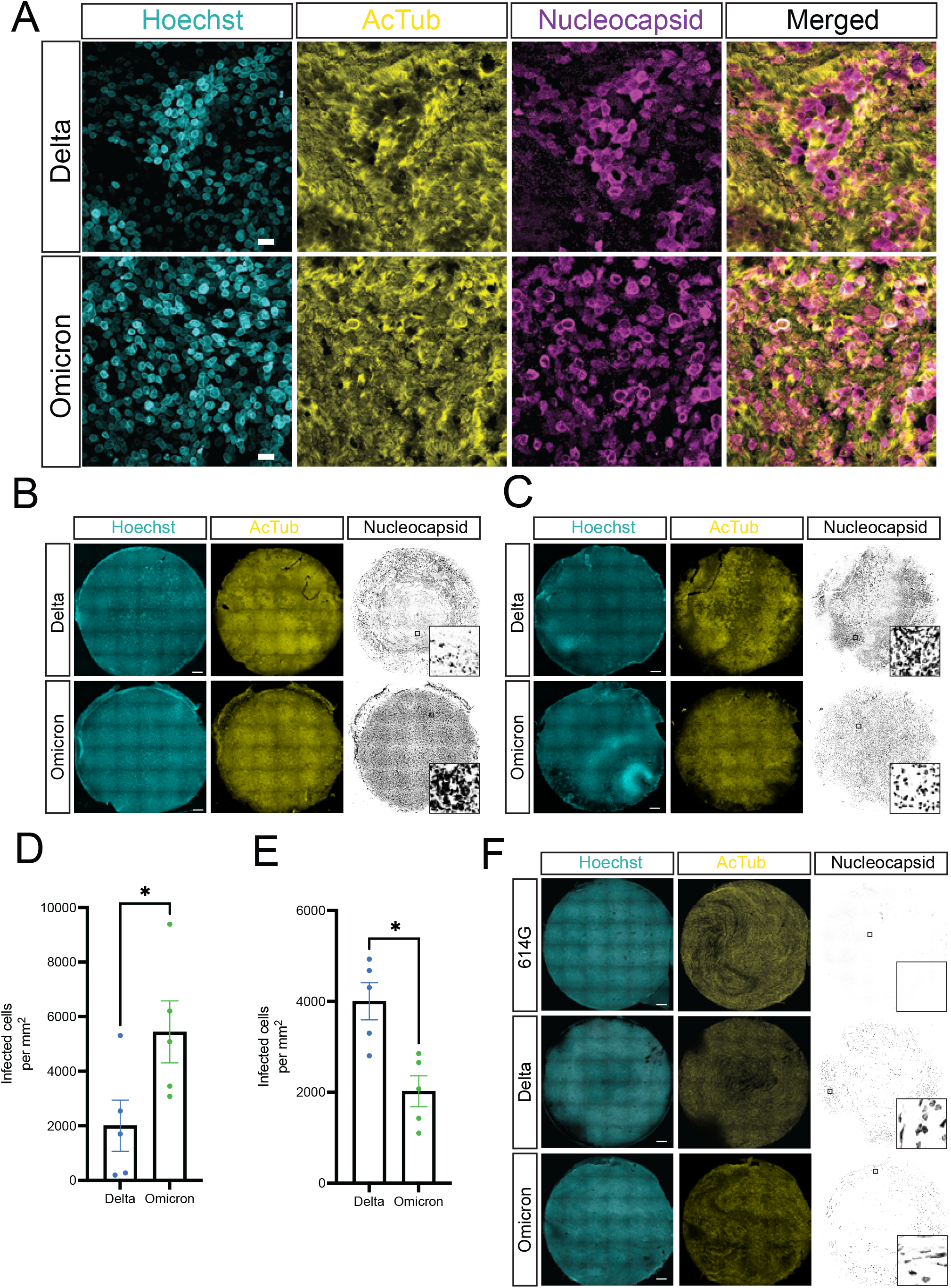
Tropism and confocal imaging of Omicron and Delta in air-liquid interface airway organoids. (a) Immunofluorescent staining of Delta and Omicron-infected 2D airway organoids at 3 d.p.i depicting infection of predominantly ciliated cells. (b-c) Whole well confocal images of Delta and Omicron-infected 2D airway organoids at 2 (b) and 3 (c) d.p.i. (d-e) Quantification of Delta and Omicron-infected cells in 2D airway organoids at 2 (d) and 3 (e) d.p.i. Groups were compared by t-test (*p < 0.05). Error bars depict SEM. (f) Whole well confocal images of 614G, Delta and Omicron-infected 2D airway organoids at 10 d.p.i. d.p.i = days post-infection. Scale bars in a represent 20 μm and scale bars in b, c and f represent 500 μm.

### Omicron rapidly outcompetes 614G, and has a fitness advantage over Delta in the first days of the infection

Next, we investigated which VOCs had the highest fitness in airway cells. For this purpose, we infected airway organoids with a mixture of 614G, Alpha, Delta or Omicron at a ratio of 1:1 or 5:1 (cumulative moi = 0.1). We sampled the cultures daily for 10 days and determined the relative frequency of both viruses to calculate replicative fitness using Sanger sequencing and electropherogram peak analyses (*32*) (**Fig. 3A**). These experiments clearly indicated that both the Alpha variant (**Fig. 3B-C, Fig. S2A-B**) and the Delta variant (**Fig. 3D-E, Fig. S2C-D**) had an increased replicative fitness in the human airway. In addition, we found that there was a minimal fitness difference between Alpha and Delta, slightly in favor of the latter (**Fig. S2E-F**), suggesting that Delta’s antigenic drift significantly contributed to its transmissibility.

**Fig. 3.**
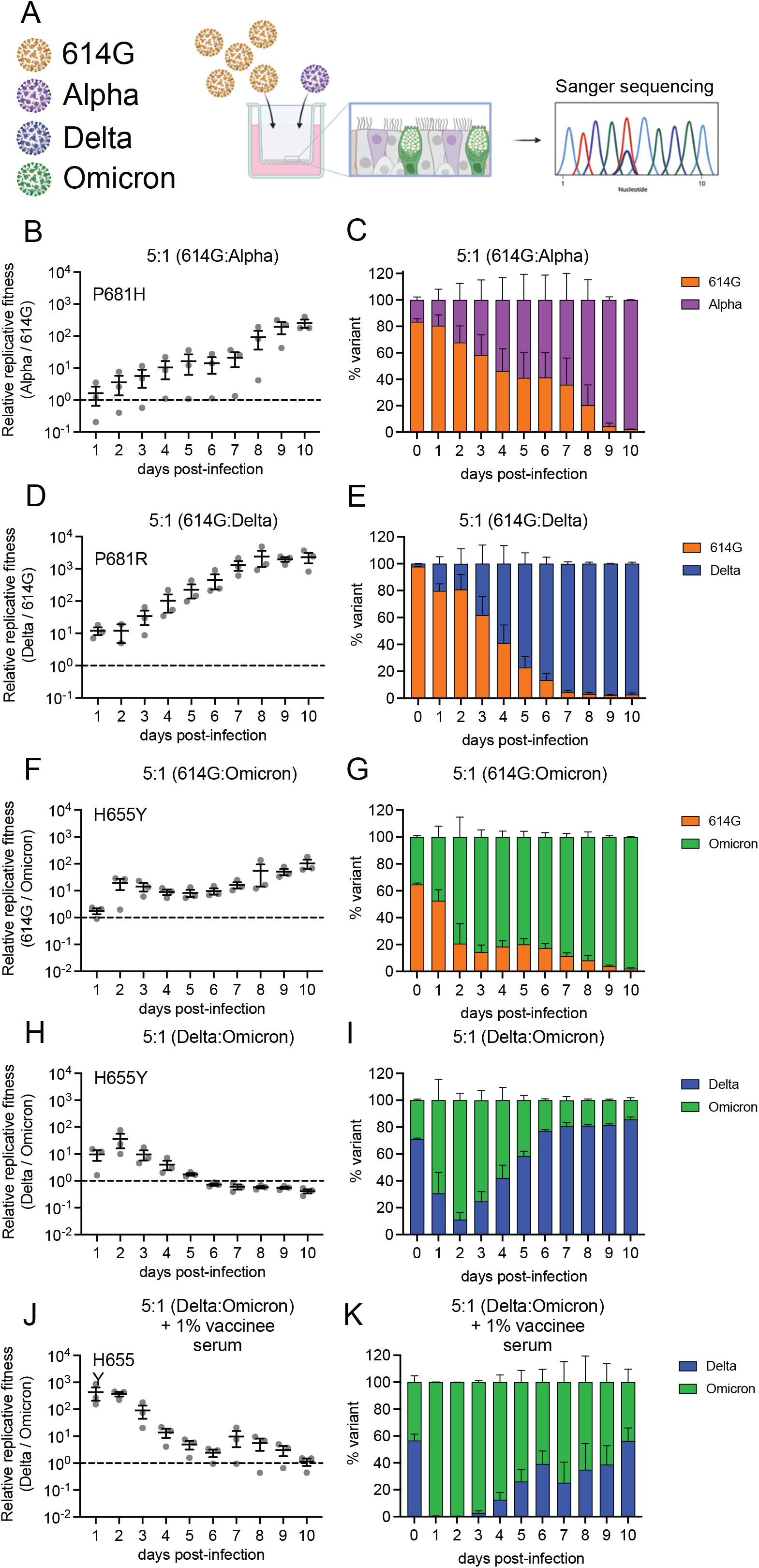
Enhanced fitness of Omicron in air-liquid interface airway organoids. (a) Graphical depiction of competition assay. (b-c) Relative replicative fitness (b) and proportion of each variant (c) in 2D airway-organoids infected with a 5:1 ratio of an ancestral 614G virus to Alpha. (d-e) Relative replicative fitness (d) and proportion of each variant (e) in 2D airway-organoids infected with a 5:1 ratio of an ancestral 614G virus to Delta. (f-g) Relative replicative fitness (f) and proportion of each variant (g) in 2D airway-organoids infected with a 5:1 ratio of an ancestral 614G virus to Omicron. (h-i) Relative replicative fitness (h) and proportion of each variant (i) in 2D airway-organoids infected with a 5:1 ratio Delta to Omicron. (j-k) Relative replicative fitness (j) and proportion of each variant (k) in 2D airway-organoids refreshed daily in the basal compartment with 1% vaccinee serum, and infected with a 5:1 ratio of Delta to Omicron. Error bars depict SEM. Panel (a) was created with BioRender.com.

Omicron has outcompeted Delta in many countries within weeks after the initial introduction, indicating that it has a high fitness (*10, 14-16*). Omicron rapidly outcompeted 614G (**Fig. 3F-G, Fig. S2G-H**). Next, we tested whether Omicron could outcompete Delta. Interestingly, Omicron had a higher fitness than Delta in the first 5 days of the infection, after which Delta outcompeted Omicron (**Fig. 2G-H, Fig. S2I-J**). These data again confirm that Omicron replicates more rapidly in the initial days of the infection (**Fig. 1F-G; Fig. 2B-E**) and that both viruses are phenotypically distinct. As infected individuals are most infectious in the first days of the infection(*29, 33, 34*), these data may explain why Omicron has rapidly outcompeted Delta in many countries. In addition, the infectious period is likely shortened by neutralizing antibodies induced upon (re-)infection or vaccination (*35, 36*). To test whether antibodies induced upon vaccination could affect the fitness of Omicron, we added serum (1%) from a vaccinated individual (2x mRNA-1273) in the lower compartment of the air-liquid interface 2D organoid wells, potentially allowing IgA antibodies to be transferred to the top compartment. Medium with serum was refreshed daily. This serum effectively neutralized Delta (IC50:2657), but had a low activity against Omicron (IC50:123.5), which is in line with recent findings on vaccine recipient serum responses against Omicron (*11, 17, 21, 22, 25, 37*). In this system, Omicron’s fitness increased and completely outcompeted Delta on day 1-3 (**Fig. 3J-K**). In the following days Omicron remained at a higher frequency until day 9, but Delta did come up slowly from day 4-10. These findings indicate that Omicron has an initial fitness advantage over Delta, and that this advantage is likely increased due to (population) immunity. They also indicate that Delta and Omicron may have evolved different strategies to evade neutralizing antibodies.

### Omicron does not efficiently spread through cell-cell fusion

SARS-CoV-2 can spread from cell to cell by fusion of infected cells with neighboring cells using the S protein, leading to the formation of large multi-nucleated syncytial cells (*38, 39*). The relevance of cell-cell spread is still being investigated, but it has been linked to antibody evasion, virus dissemination and pathogenicity (*40, 41*). As Delta could still outcompete Omicron late in infection in the presence of neutralizing serum with a high anti-Delta activity (**Fig. 3J-K**), we hypothesized that Delta may spread more efficiently through cell-cell fusion than Omicron. In airway organoids, we noticed that Delta resulted in abundant syncytial cells, whereas Omicron did not (**Fig. 4A-B**), showing that indeed Delta is more fusogenic. Using a GFP-complementation system and S protein overexpression (*42*) (**Fig. 4C**) we observed that the Delta S was more fusogenic compared to 614G in VeroE6, VeroE6-TMPRSS2 and Calu-3 cells (**Fig. 4D-F**). The Omicron S was significantly less fusogenic in all cells. Next, we investigated whether similar results would be obtained using an authentic virus fusion assay, also based on GFP complementation in Calu-3 cells (**Fig. 4G**). In this assay, Delta was again more fusogenic, whereas infection with Omicron did not result in any detectable fusion. Our findings in organoids and cell lines indicate that Omicron does not spread efficiently through cell-cell fusion. As S-mediated fusogenicity is directed by the expression of S-activating proteases on the cell surface (*43, 44*) and Omicron contains mutations close to the S1/S2 cleavage site, we hypothesized that the Omicron S may have evolved to use other proteases.

**Fig. 4.**
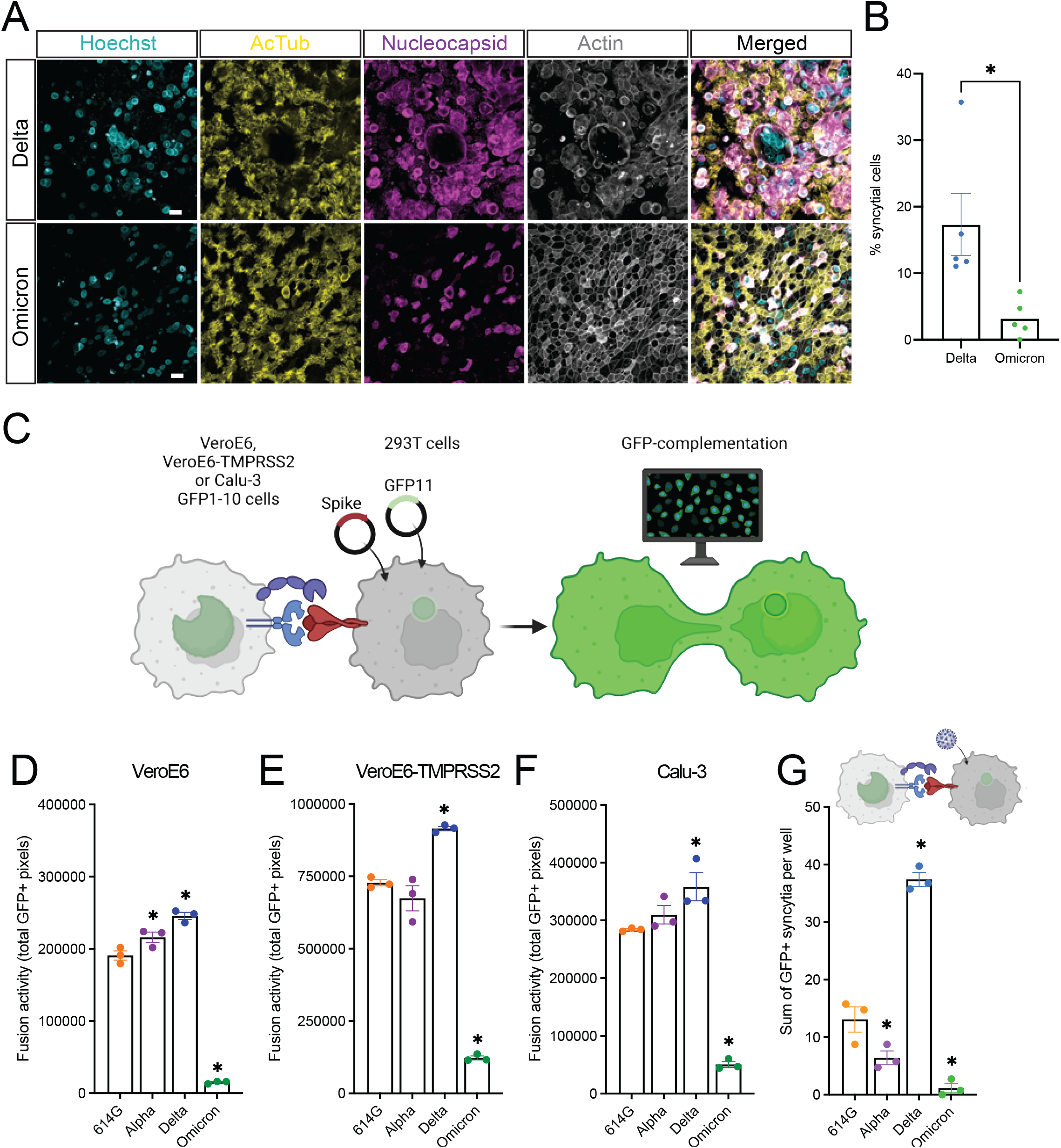
Omicron spreads less efficiently through cell-cell fusion compared with Delta. (a) Human 2D airway organoids infected with Omicron and Delta at 3 d.p.i were stained for nuclei (hoechst; blue), cilia (AcTub; yellow), actin (phalloidin; grey) and SARS-CoV-2 nucleocapsid (purple) to image syncytial cell formation. (b) Quantification of (a) by manual counting of syncytial cells per field, represented as % of total infected cells per field. A syncytial cell was defined as a cell with positive nucleoprotein staining, two or more nuclei that are not separated by actin rings. Data is depicted as mean with SEM. Statistical analysis was performed by t-test. (c) Graphical depiction of the GFP-complementation fusion assay. (d-f) Quantification of 614G, Alpha, Delta and Omicron S-mediated cell-cell fusion by measuring the sum of all GFP+ pixels per well on VeroE6, VeroE6-TMPRSS2 and Calu-3 cells 18 hours post transfer. (g) Quantification of 614G, Alpha, Delta and Omicron authentic virus GFP-complementation fusion assay on Calu-3 cells by counting the sum of all GFP+ syncytia per well at 24 h p.i. Error bars depict SEM. Groups in d-g were compared by one-way ANOVA comparing VOCs to 614G. *p < 0.05.

### Omicron is less dependent on TMPRSS2 for entry and has evolved to use an alternative protease in airway cells

SARS-CoV-2 S is proteolytically cleaved to activate the S protein for fusion. This occurs in producing cells at the S1/S2 site by furin, and at a site directly adjacent to the fusion peptide by TMPRSS2 at the cell surface or cathepsins in the endosome (*45, 46*). Coronavirus tropism is mainly influenced by receptor and protease usage (*43, 44*). Therefore, we first investigated whether Omicron uses TMPRSS2 for entry. For this purpose, we used a S-based VSV pseudovirus system described before (*42*) (**Fig. 5A**). Omicron was 1-2 log less infectious on VeroE6 cells and 2-3 log less infectious on VeroE6-TMPRSS2 cells (**Fig. 5B**). In line with previous data (*42, 47*), TMPRSS2 expression effectively increased the infectivity of 614G, Alpha and Delta pseudoviruses (**Fig. 5B**), but surprisingly Omicron’s infectivity was not affected by TMPRSS2. We confirmed this data by investigating the route of entry of Omicron pseudoviruses in cells that have an active TMPRSS2 as well as a cathepsin-mediated entry route (*42*). TMPRSS2-mediated entry was inhibited using serine protease inhibitor camostat and cathepsins were inhibited using E64D (**Fig. 5C-D**). Whereas 614G, Alpha, and Delta entered these cells exclusively through TMPRSS2, Omicron used this entry route inefficiently, leading to some endosomal entry, and confirming that Omicron does not efficiently use TMPRSS2. Next, we determined the infectivity of Omicron on Calu-3 cells, in which SARS-CoV-2 generally enters through TMPRSS2 (*42, 48*). These cells lack a cathepsin-mediated entry route (*42, 49*). Omicron was ∼1 log less infectious on Calu-3 cells (**Fig. 5E**), but was still inhibited by camostat and not E64D (**Fig. 5F-G**). These data indicate that entry into Calu-3 cells occurs either inefficiently through TMPRSS2, or is mediated by another serine protease that is blocked by camostat. As we noticed that the S1/S2 cleavage of Omicron pseudovirus (produced on 293T cells) was much lower than authentic virus (produced on Calu-3 cells) (**Fig. S3A-D**), we used authentic viruses to verify that Omicron did not benefit from TMPRSS2 and had a lower infectivity on Calu-3 cells (**Fig. 5H-I**). Although in this case, pseudovirus and authentic virus data led to the same conclusion, these data indicate that pseudovirus data should be confirmed using authentic viruses. Next, we tested which entry route is used by Omicron and Delta authentic viruses on airway organoids. Both viruses were effectively inhibited by camostat (10 μM), and not E64D (10 μM), or the combination of both (**Fig. 5J-K**), indicating that both viruses use the serine protease and not the cathepsin-mediated entry pathway in relevant cells. These findings were reproduced in a second experiment using a higher camostat concentration (100 μM) (**Fig. S4A-B**). In line with previous work (*42*), E64D was effective against all viruses in VeroE6 cells (**Fig. S5A-B**). In addition, these findings, together with the observation that Omicron replicates more efficiently in airway cells compared with Delta and does not efficiently use TMPRSS2, suggest that Omicron has evolved to use another serine protease expressed by airway cells.

**Fig. 5.**
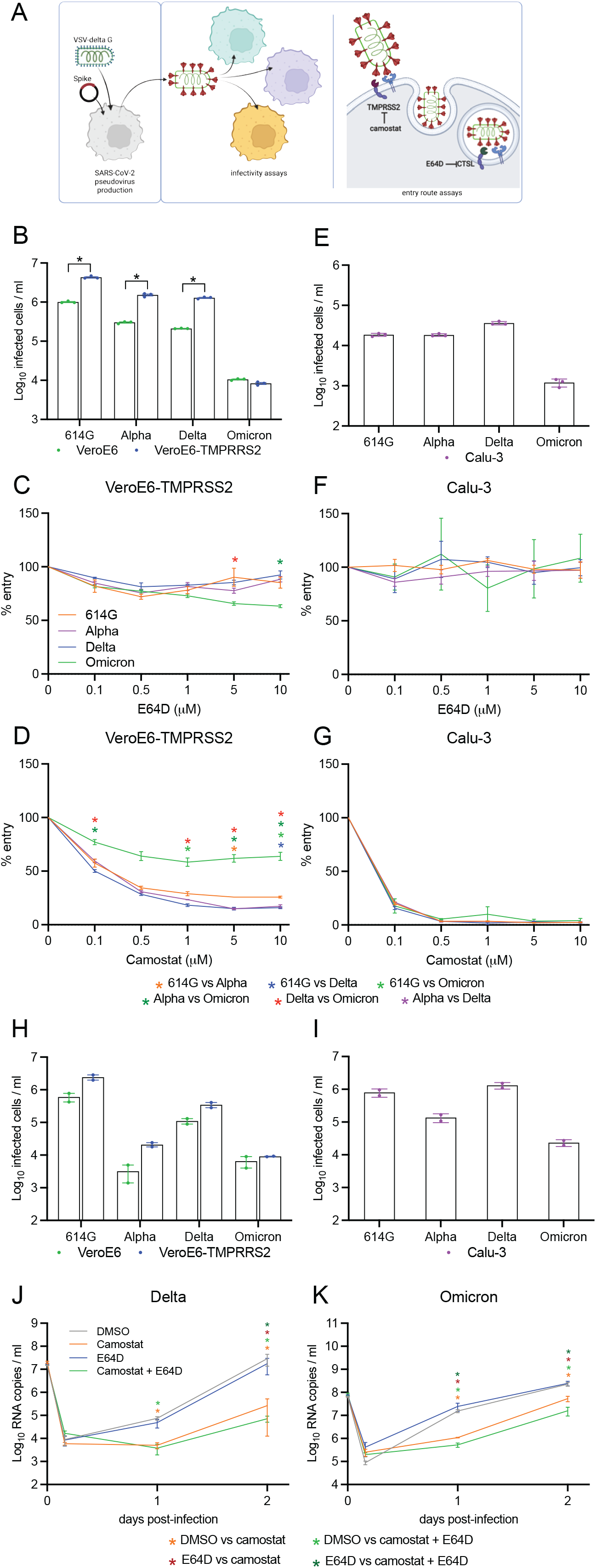
Omicron does not efficiently enter cells using TMPRSS2. (a) Graphical depiction of VSV pseudovirus production and subsequent infectivity and entry route assays. (b) Infectivity of pseudoviruses in VeroE6 and VeroE6-TMPRSS2 cells. (c-d) Percentage entry of pseudoviruses in VeroE6-TMPRSS2 cells pretreated with a concentration range of either E64D (c) or camostat (d). (e) Infectivity of pseudoviruses in Calu-3 cells. (f-g) Percentage entry of pseudoviruses in Calu-3 cells pretreated with a concentration range of either E64D (f) or camostat (g). (h-i) Infectivity of authentic viruses in VeroE6, VeroE6-TMPRSS2 (h), and Calu-3 (i) cells. (j-k) Replication of Delta (j) and Omicron (k) in the presence of DMSO (vehicle), camostat (10 μM), E64D (10 μM), or the combination of camostat and E64D in human airway organoids. Groups were compared by two-way ANOVA. *p < 0.05. Error bars depict SEM. Panel (a) was created with BioRender.com.

### Omicron does not efficiently infect AT2 cells

The ability of SARS-CoV-2 to infect alveolar type 2 (AT2) cells is thought to be an important virulence factor for SARS-CoV-2, but also SARS-CoV and MERS-CoV, which depend on TMPRSS2 for entry (*46*). Inflammation in the alveoli likely sets off an inflammatory response ultimately leading to flooding of the alveoli, impeding gas exchange. AT2 cells express TMPRSS2 and removing TMPRSS2 activity effectively reduces pathology in animals for SARS-CoV, SARS-CoV-2 and H7N9 influenza A (*50, 51*). Until recently studying SARS-CoV-2 pathogenesis in these cells has been difficult due to the lack of an *in vitro* model, but recently several groups have established organoid models of the human adult alveolus (*52-55*). As Omicron does not efficiently use TMPRSS2 and preliminary reports have indicated that Omicron might be less pathogenic (*56-60*), we used an alveolar organoid model (adapted from (*52, 53*)) (**Fig. S6**) to assess whether Omicron could infect AT2 cells (**Fig. 6A-C**). Whereas Delta efficiently replicated in AT2 cultures, Omicron was severely attenuated (**Fig. 6D-E**). These observations were confirmed in AT2 organoids from another donor (**Fig. 6F-G**) and supported by confocal microscopy at day 4 post-infection, which showed abundant infection for Delta, but not for Omicron (**Fig. 6H-I**). In line some infectious virus production late in the infection for Omicron, we did observe infected cells in Omicron infected AT2 organoids at day 10 post-infection (**Fig. S7A-B**). Altogether, our data indicate that Omicron does not efficiently infect AT2 cells, whereas airway infection is highly efficient (**Fig. 1-3**).

**Fig. 6.**
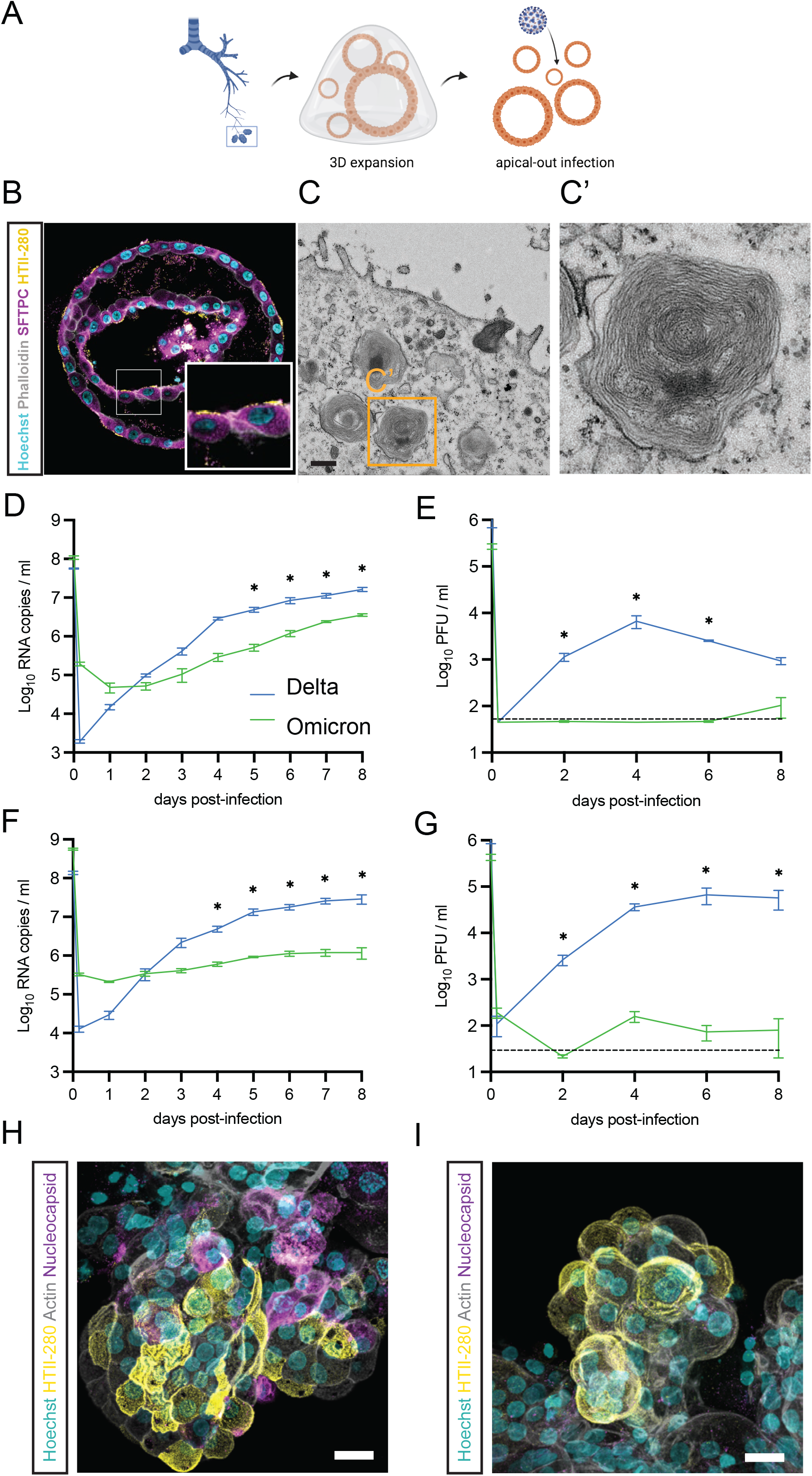
Omicron does not efficiently infect AT2 cells. (a) Graphical depiction of AT2 cell isolation, 3D expansion and apical-out infection. (b) Immunofluorescent staining of 3D AT2 cells for AT2 cell markers surfactant protein C (SFTPC; purple) and HTII-280 (yellow). (c-c’) Transmission electron microscopy images of 3D AT2 cells depicting surfactant-producing lamellar bodies. (d-g) Replication kinetics of Omicron and Delta in two donors (d-e, donor 1; f-g, donor 2) of apical-out 3D AT2 cells. Donor 1 was sorted, whereas donor 2 was passaged to generate AT2 organoids. Replication kinetics were quantified by RNA copies (d and f) and PFU (e and g) per ml. (h-i) Immunofluorescent images of Delta (h) and Omicron (i) infected 3D AT2 cells at 4 d.p.i. Scale bars in h-I represent 20 μm. Groups were compared by two-way ANOVA. *p < 0.05. Error bars depict SEM. Panel (a) was created with BioRender.com.

After nearly two years of circulation and evolution of SARS-CoV-2 in humans, the highly mutated Omicron variant emerged, raising concerns about its potential extensive immune evasion, increased transmissibility and pathogenicity. Here, we used human organoid systems to investigate Omicron’s shedding, fitness and replicative potential in the airways and alveoli. We report that Omicron efficiently replicates ciliated cells of the airway and has a higher fitness than 614G and Delta. Mechanistically, Omicron was much less fusogenic and did not efficiently use TMPRSS2 for entry. However, Omicron was inhibited by serine protease inhibitor camostat (and not cathepsin inhibitor E64D) in airway cells indicating that it has evolved to use another serine protease. Our observation that Omicron does not efficiently replicate in AT2 cells suggests that this protease is not present on these cells. Our data are in line with preliminary reports indicating that Omicron’s hospitalization rates appear lower than Delta’s (*56-60*). At the same time, our data on Omicron’s rapid replication and competitive fitness in the airway support epidemiological observations that Omicron appears to be even more transmissible than Delta (*14, 15, 61, 62*). A recent study in hamsters showed that Omicron was attenuated in both the lower and upper respiratory tract (*63*). Although these data support that Omicron may be less pathogenic, they fail to explain why Omicron spreads so efficiently in humans. Our data imply that human organoid models are well suited for studying viral fitness and pathogenicity, especially when comparing viral variants that are adapting to specific human host factors.

## Acknowledgements

We thank Robbert Rottier for providing human adult lung material, Pim French for technical support, and the Microscopy CORE lab at Maastricht University for the TEM sample preparation.

## Author contributions

Conceptualization: M.M.L., A.Z.M., T.I.B., B.L.H.

Methodology: M.M.L., A.Z.M., T.I.B, N.C.W., P.J.P., M.J.M., M.W., M.R.

Investigation: M.M.L., A.Z.M., T.I.B, J.Z., Y.W., N.G., K.K., D.S., P.B.D., R.V.A., T.B., C.H.G., M.W., M.R.

Visualization: M.M.L., A.Z.M., T.I.B.

Funding acquisition: B.L.H.

Supervision: B.L.H.

Writing – original draft: M.M.L., A.Z.M., T.I.B.

Writing – review & editing: B.L.H.

## Supplementary Materials

### Material and Methods

#### Cell lines

Vero and VeroE6 cells were maintained in Dulbecco’s modified Eagle’s medium (DMEM, Gibco) supplemented with 10% fetal bovine serum (FBS), HEPES (20 mM, Lonza), sodium pyruvate (1 mM, Gibco), penicillin (100 IU/mL), and streptomycin (100 IU/mL) at 37°C in a humidified CO_2_ incubator. Calu-3 cells were maintained in Opti-MEM I (1) + GlutaMAX (Gibco) supplemented with 10% FBS, penicillin (100 IU/mL), and streptomycin (100 IU/mL) at 37°C in a humidified CO_2_ incubator. HEK-293T cells were cultured in DMEM supplemented with 10% FCS, sodium pyruvate (1 mM, Gibco), non-essential amino acids (1×, Lonza), penicillin (100 IU/mL), and streptomycin (100 IU/mL) at 37°C in a humidified CO2 incubator. TMPRSS2, GFP11 and GFP1-10 stable cells were maintained in a medium containing hygromycin (Invitrogen), puromycin (Invivogen) and geneticin (Invitrogen), respectively. Cell lines were tested negative for mycoplasma.

#### Plasmids

Codon-optimized SARS-CoV Wuhan-D614G, Alpha and Delta S expression plasmids (pLV) were ordered from Invivogen. The pGAGGS-Omicron expression plasmid was kindly provided by Dr. Berend Jan Bosch. All S expressing plasmids contained a deletion of the last 19 amino acids containing the Golgi retention signal of the SARS-CoV S protein. The plasmids were used for S pseudovirus production and the GFP-complementation fusion assay. The pGAGGS-β-Actin-P2A-7xGFP11-BFP plasmid was cloned into pQCXIP and used for retrovirus production and subsequent generation of the GFP11 stable cells as previously described by Mykytyn et al. (*42*).

#### Airway cell isolation, culture and differentiation

Human adult airway stem cells were isolated from bronchus (n=1) and lung parenchyma (n=3) as described previously (*28*) using a protocol adapted from Sachs and colleagues (*64*). Cells were differentiated using commercially available Pneumacult-ALI medium (complete base medium with 1X maintenance supplement; Stemcell) as described before(*26, 28*). Cells were differentiated at the air-liquid interface for 3-4 weeks. Medium was replaced every 4-5 days.

Adult human lung tissue was obtained from non-tumour lung tissue obtained from patients undergoing lung resection. Lung tissue was obtained from residual, tumor-free, material obtained at lung resection surgery for lung cancer. The Medical Ethical Committee of the Erasmus MC Rotterdam granted permission for this study (METC 2012-512).

Bronchus derived organoids were used in Figure 1C-F and Figure S2E-F and lung parenchyma derived organoids were used in Figure 1A-B, Figure 2, Figure 3, Figure 5, and Figure S2A-D, G-J, S4.

#### Alveolar cell isolation, culture and differentiation

Human adult alveolar stem cells were isolated using a protocol adapted from Lamers and colleagues (2020). Distal lung pieces were chopped into 2 × 2 mm pieces, washed twice with cold PBS and incubated in 100% dispase (Corning), supplemented with 10 μM Y-27632 (MedChemExpress) for 30-60 min at 37°C. Next, cells were dissociated from the tissue by pipetting using a 5 ml serological pipet and subsequently using a P1000 micropipette. Cells were then strained using a 100 μm cell strainer (Falcon) and washed twice in cold AdDF+++. Red blood cells were lysed using red blood cell lysis buffer (Roche), after which cells were again washed twice with cold AdDF+++. Cells were pelleted, incubated on ice for 2 min, and plated in growth factor reduced basement membrane extract (BME; Cultrex). After BME solidification, alveolar medium (recipe (see below) adapted from Youk and colleagues (*53*) was added and cells were incubated at 37°C in a humidified CO_2_ incubator (Youk et al., 2020). Alveolar medium consisted of AdDF+++ containing B27 (1X; minus vitamin A; Invitrogen), N2 (1X; Invitrogen), *N*-acetylcysteine (1.25 mM), CHIR99021 (3 μM; Sigma), RSPO1 (R-spondin 1; conditioned medium, 10% v/v), FGF7 (100 ng/ml; Peprotec), FGF10 (100 ng/ml; Peprotec), epidermal growth factor (EGF; 50 ng/ml; Peprotec), NOG (NOGGIN; 100 ng/ml; Peprotec), SB431542 (10 μM; Tocris), Neuregulin-1 (100 ng/ml; Peprotec)(shown to increase the growth of alveolar organoids (*52*)), and Primocin (1X; Invivogen). Y-27632 (10 μM) was added for the first 5 days. Medium was replaced every 4-5 days. The Medical Ethical Committee of the Erasmus MC Rotterdam granted permission for this study (METC 2012-512).

To obtain pure AT2 organoids, organoid lines were either passaged in alveolar medium for 4 passages or sorted. Sorted organoids were used in Figure 6B-E, H, I, Figure S6, S7, and unsorted organoids from another donor were used in Figure 6F-G.

For sorting, organoids were digested using TrypLE Express and washed in AdDF++ twice before incubating the cells in Lysotracker (Thermofisher) for 20 min at 37°C. Next, cells were incubated in FACS buffer (2mM EDTA, 2.5% bovine serum albumin (BSA) in PBS) on ice for 5 min, stained with the AT2 marker antibody HTII-280 (1:40; Terrace Biotech) on ice for 15 min, and with goat anti-mouse IgM Alexa Fluor 488 (1:400; Invitrogen) for 5 min. Cells were then washed once in FACS buffer and HTII-280-high and Lysotracker-high cells were sorted using a FACS Aria cell sorter (BD biosciences) into AdDF+++ containing 10 μM Y-27632. Cells were pelleted, incubated on ice for 2 min and plated in BME domes in alveolar medium. Y-27632 (10 μM) was added for the first 5 days. Cells were incubated at 37°C in a humidified CO_2_ incubator and medium was replaced every 4-5 days.

Sorted and unsorted AT2 organoids were split every 2-3 weeks at a 1:3-1:8 ratio. Organoids were dissociated to small clumps using TrypLE.

All centrifugation steps were performed at 400 xg for 3 min.

#### Viruses

SARS-CoV-2 isolates were grown to passage 3 on Calu-3 (ATCC HTB-55) cells in Advanced DMEM/F12 (Gibco), supplemented with HEPES, Glutamax, penicillin (100 IU/mL) and streptomycin (100 IU/mL) at 37°C in a humidified CO_2_ incubator. Infections were performed at a multiplicity of infection (moi) of 0.01 and virus was harvested after 72 hours. The culture supernatant was cleared by centrifugation at 1000 x g for 5 min and stored at −80°C in aliquots.

For plaque assay titrations, virus stocks or experimental samples were thawed and diluted in 10-fold serial dilutions in 200μl Opti-MEM I (1X) + GlutaMAX. 100μl of each dilution was added to monolayers of 1 × 10^6^ VeroE6, VeroE6-TMPRSS2 or Calu-3 cells in the same medium in a 12-well plate. Cells were incubated with inoculums at 37°C for 4 hours and then medium was replaced for 1.2% Avicel (FMC biopolymers) in Opti-MEM I (1X) + GlutaMAX for two days. To determine the 8-hour infectious titer, virus stocks were thawed and diluted in 10-fold serial dilution in 200μl Opti-MEM I (1X) + GlutaMAX. 100μl of each dilution was added to monolayers of 1 × 10^5^ VeroE6, VeroE6-TMPRSS2 or Calu-3 cells in a 96-well plate. Cells were incubated with inoculums at 37°C for 8 hours. Cells were fixed in 4% formalin for 20 minutes, permeabilized in 70% ice-cold ethanol and washed in PBS. Cells were blocked in 0.6% BSA (bovine serum albumin; Sigma) in PBS and stained with rabbit anti-nucleocapsid (Sino biological; 1:2000) in PBS containing 0.6% BSA, washed thrice in PBS, and stained with goat anti-rabbit Alexa Fluor 488 (Invitrogen; 1:4000) in PBS containing 0.6% BSA. Cells were washed twice in PBS and plates were scanned on the Amersham Typhoon Biomolecular Imager (channel Cy2; resolution 10 μm; GE Healthcare). All staining steps were performed at room temperature for one hour. Plaque assay analysis was performed using ImageQuant TL 8.2 software (GE Healthcare). All work with infectious SARS-CoV-2 was performed in a Class II Biosafety Cabinet under BSL-3 conditions at Erasmus Medical Center.

#### SARS-CoV-2 infections

Prior to SARS-CoV-2 infection of 2D and 3D cultures, cells were washed three times. Washes were performed apically for 2D airway organoid cultures with 300 μl AdDF+++ medium. To assess the effect of camostat mesylate (Sigma) and E64D (MedChemExpress) on viral replication, 2D cultures were pretreated in the basal and apical compartment for 1 hour with 10μM of either compound or the combination of both, or DMSO in the negative control Submerged 2D monolayers of VeroE6 and Calu-3 cells were washed in 300 μl PBS and subsequent infection was performed in AdDF +++ medium. 3D AT2 cultures were incubated with TrypLE Express for 5 minutes at room temperature and washed three times with 5 ml AdDF+++ medium. Cells were incubated with inoculums for 2-4 hours at 37°C in a humidified CO_2_ incubator at a moi of 0.1. Washing was performed three times at 4 hours post infection in AdDF+++ after which:

- For 2D ALI cultures, a fourth wash was performed and collected as the 4 hours post-infection baseline.
- For 2D submerged VeroE6 and Calu-3 cell cultures, 500 μl AdDF +++ was added to the wells and incubated for 10 minutes before a sample was taken from the supernatant as the 4 hours post-infection baseline.
- For 3D AT2 cultures, organoids were plated in a slurry of 5% BME in 800 μl alveolar medium per well in 24 well plates. In this slurry most organoids adopted an apical-out orientation. A baseline sample was taken 1 hour after plating to first allow organoids to settle within the slurry.

For 2D ALI cultures, samples were taken at the indicated time points as follows: 500 μl for 12mm inserts or 200μl for 6.5mm inserts of AdDF+++ was added to the apical side of the cells and cells were incubated for 10 min at 37°C in a humidified CO_2_ incubator after which supernatants were pipetted up and down on the cells twice and transferred to a microvial. After collecting, for experiments with camostat mesylate or E64D, the basal compartment medium was replaced daily with medium containing fresh compound. For 2D submerged cultures, 150 μl samples were taken directly from the supernatant for storage in - 80°C and 60 μl was transferred directly to 90 μl MagnaPure LC Lysis buffer (Roche) for RNA isolations. 210 μl fresh medium was added to the cultures. For 3D AT2 cultures, 60 μl supernatant samples were taken directly from the supernatant for storage in -80°C and 30 μl was transferred directly to 45 μl MagnaPure LC Lysis buffer for RNA isolations. 90 μl fresh expansion medium was added to the cultures. All supernatant samples were stored at -80°C until further processing for viral titer determinations. All replication curves were performed in triplicate.

#### Determination of virus titers using qRT-PCR

SARS-CoV-2 RNA was extracted as described previously (*26*)and RNA genome copies (E-gene) were determined by qRT-PCR. Briefly, supernatant samples were thawed and centrifuged at 2,000 × g (supernatant) for 3 min. 60 μl sample was lysed in 90 μl MagnaPure LC Lysis buffer (Roche) at RT for 10 min. RNA was extracted by incubating samples with 50 μl Agencourt AMPure XP beads (Beckman Coulter) for 15 min at room temperature, washing beads twice with 70% ethanol on a DynaMag-96 magnet (Invitrogen) and eluting in 30 μl DEPC-treated water. RNA copies per ml were determined by qRT-PCR using primers targeting the E gene and comparing the Ct values to a counted standard curve derived from a 614G stock.

#### Immunofluorescent staining

Transwell inserts with 2D airway organoids were fixed in 4% formalin, permeabilized in 70% ethanol and blocked for 60 min in blocking buffer (10% filtered normal goat serum (MP Biomedicals) in PBS. Cells were incubated with primary antibodies overnight at 4°C in blocking buffer, washed twice with PBS, incubated with corresponding secondary antibodies (Alexa 488- / 594-conjugated anti-rabbit IgG, anti-mouse IgG, or anti-mouse IgG1 (1:500; Invitrogen)) in blocking buffer for 2 hours at room temperature, washed two times with PBS, incubated with indicated additional stains (hoechst (ThermoFisher) or phalloidin (Santa Cruz)), washed twice with PBS, and mounted in Prolong Antifade (Invitrogen) mounting medium. Viral nucleocapsid was stained with rabbit anti-NP (1:500; Sino biological), ciliated cells were stained with anti-AcTub (1:100; Santa Cruz, 6-11B-1).

3D Alveolar organoids were fixed in 4% formalin, permeabilized in 0.1% Triton X-100, and blocked for 60 min in 10% normal goat serum in PBS (blocking buffer). Cells were incubated with primary antibodies overnight at 4°C in blocking buffer, washed twice with PBS, incubated with corresponding secondary antibodies (Alexa488- and 594-conjugated anti-rabbit IgG, anti-mouse IgG, or anti-mouse IgG1 (1:500; Invitrogen)) in blocking buffer for 2 hours at room temperature, washed two times with PBS, incubated with indicated additional stains (hoechst or phalloidin), washed twice with PBS, and mounted in Prolong Antifade mounting medium. Viral nucleocapsid was stained with rabbit anti-NP, AT2 cells with anti-HTII-280 (1:200; Terrace biotech) or anti-SFPTC (1/200; Millipore) and basal cells with anti-TP63.

All confocal imaging was performed on a LSM700 confocal microscope using ZEN software.

#### Replicative fitness measurements

Replicative fitness was measured as described before (*32*). Briefly, air-liquid airway organoids differentiated in Pneumacult were infected as described above at a 1:1 or 5:1 ratio at a final moi of 0.1 (based of PFU/ml) in triplicate. Apical washes were performed daily and stored at - 80°C until further processing. RNA was extracted as described above and samples were subjected to cDNA synthesis using Superscript IV (Invitrogen), according to the manufacturer’s instructions. Next, a ∼700 bp region in the S gene containing the P681H/R and H655Y mutations was amplified by PCR using Pfu UltraII (Agilent) using the following primers:

Forward: ‘5-TGA CAC TAC TGA TGC TGT CCG TG-3’

Reverse: ‘5-GAT GGA TCT GGT AAT ATT TGT GG-3’

Conditions for the PCR were as follows:

1. Denaturation for 5 minutes at 94°C;
2. Denaturation for 20 seconds at 94°C;
3. Annealing for 20 seconds at 52°C;
4. Extension for 60 seconds at 72°C;
5. Steps 2-4 40 times;
6. Final extension for 5 minutes at 72°C.

Next, amplified PCR products were run on agarose gel to verify successful amplification and the remaining PCR products were purified using a PCR purification kit (Qiagen) before Sanger sequencing using the forward primer. Sanger sequences were analyzed using QSVanalyser (Insilicase). Raw peak heights were calculated of the variant-specific polymorphism at position 681 or 655 of spike. Replicative fitness was calculated according to w = (f0/i0), in which i0 is the initial peak height ratio and f0 is the ratio at a given time point (Plante et al., 2021a).

#### Deep-sequencing

Viral genome sequences were determined using Illumina deep-sequencing. RNA was extracted as described above and cDNA was generated using ProtoscriptII reverse transcriptase enzyme (New England, BiotechnologyBioLabs) according to the manufacturer’s protocol. Samples were amplified using the QIAseq SARS-CoV-2 Primer Panel (Qiagen). Amplicons were purified with 0.8× AMPure XP beads and 100ng of DNA was converted to paired-end Illumina sequencing libraries using the KAPA HyperPlus library preparation kit (Roche) with the KAPA unique dual-indexed adapters (Roche) as per manufacturers recommendations. The barcode-labeled samples were pooled and analyzed on an Illumina sequencer V3 MiSeq flowcell (2×300 cycles). Sequences were analyzed using CLC Genomics Workbench 21.0.3. The 614G virus (clade B; isolate Bavpat-1; European Virus Archive Global #026 V-03883) passage 3 sequence was identical to the passage 1 (kindly provided by Dr. Christian Drosten) and no minor variants >20% were detected. No other minor variants >20% were detected. The 614D, Delta (clade B.1.617.2) and Omicron (clade B.1.1.529, BA-1) variant passage 3 sequences were identical to the original respiratory specimens and no minor variants >20% were detected. Due to primer mismatches in the S1 region of the omicron spike gene, amplicons 72, 73, 75 and 76 were sequenced at low coverage. Therefore, the S1 regions of the original respiratory specimen and passage 3 virus were confirmed to be identical by Sanger sequencing. The Delta and Omicron sequences have been submitted to Genbank.

The Delta variant contained the following spike changes: T19R, G142D, del156-157, R158G, A222V, L452R, T478K, D614G, P681R and D950N. The Omicron variant contained the following spike mutations: A67VS, del69-70, T95I, G142-, del143-144, Y145D, del211, L212I, ins215EPE, G339D, S371L, S373P, S375F, K417N, N440K, G446S, S477N, T478K, E484A, Q493R, G496S, Q498R, N501Y, Y505H, T547K, D614G, H655Y, N679K, P681H, N764K, D796Y, N856K, Q954H, N969K, L981F.

#### Bulk mRNA sequencing

Library preparation was performed at Single Cell Discoveries (Utrecht, The Netherlands), using an adapted version of the CEL-seq protocol. In brief: total RNA was extracted using the standard TRIzol (Invitrogen) protocol and used for library preparation and sequencing. mRNA was processed as described previously, following an adapted version of the single-cell mRNA seq protocol of CEL-Seq (*65, 66*). Samples were barcoded with CEL-seq primers during a reverse transcription and pooled after second strand synthesis. The resulting cDNA was amplified with an overnight *in vitro* transcription reaction. From this amplified RNA, sequencing libraries were prepared with Illumina Truseq small RNA primers. Paired-end sequencing was performed on the Illumina NextSeq500 platform using barcoded 1 × 75 nucleotide read setup. Read 1 was used to identify the Illumina library index and CEL-Seq sample barcode. Read 2 was aligned to the CRCh38 human RefSeq transcriptome, with the addition of SARS-CoV-2 (Ref-SKU: 026V-03883; MW947280) genomes, using BWA using standard settings (*67*). Reads that mapped equally well to multiple locations were discarded. Mapping and generation of count tables was done using the MapAndGo script (https://github.com/anna-alemany/transcriptomics/tree/master/mapandgo). Data has been deposited in GEO (GSE178333). SARS-CoV-2-mapping reads were removed for further analysis. Differential expression analysis were performed using the DESeq2 package(*68*). P-values resulting from the differential expression tests were adjusted with Benjamini & Hochberg correction and considered significant with Padj<0.05 (*69*). Shrinkage is applied to the resulting fold changes. To illustrate the results, the significantly differentially regulated genes with an absolute fold change above 2 (with a maximum of 50 genes) are depicted in heatmaps, using expression values after variance stabilizing transformation.

#### Transmission electron microscopy

Organoids were chemically fixed for 3 hours at room temperature with 1.5% glutaraldehyde in 0.067 M cacodylate (pH 7.4) and 1 % sucrose. Samples were washed once with 0.1 M cacodylate (pH 7.4), 1 % sucrose and 3 times with 0.1 M cacodylate (pH 7.4), followed by incubation in 1% osmium tetroxide and 1.5% K_4_Fe(CN)_6_ in 0.1 M sodium cacodylate (pH 7.4) for 1 hour at 4°C. After rinsing with Milli-Q water, organoids were dehydrated at room temperature in a graded ethanol series (70%, 90%, up to 100%) and embedded in epon. Epon was polymerized for 48h at 60°C. Ultrathin sections (60 nm) were cut using a diamond knife (Diatome) on a Leica UC7 ultramicrotome, and transferred onto 50 Mesh copper grids covered with a formvar and carbon film. Sections were post-stained with uranyl acetate and lead citrate. All TEM data were collected autonomously as virtual nanoscopy slides (*70*) on FEI Tecnai T12 microscopes at 120kV using an Eagle camera. Data was stitched, uploaded, shared and annotated using Omero (annotated using Omero) and PathViewer.

#### Pseudovirus assays

Pseudovirus production, infectivity, and entry assays were performed as described before (*42*). Briefly, pseudoviruses were generated using spike pseudotyping vectors from Invivogen. Pseudoviruses expressing 614G, Alpha, Delta and Omicron S were titrated in triplicate by preparing 10-fold serial dilutions in Opti-MEM I (1×) + GlutaMAX (Gibco). 30 μl of each dilution was transferred to 2 × 10^4^ VeroE6 and VeroE6-TMPRSS2 cell monolayers or 8 × 10^4^ Calu-3 cell monolayers in the same medium in a 96-well plate. The cells were incubated at 37°C overnight and then scanned on the Amersham Typhoon Biomolecular Imager (channel Cy2; resolution 10 μm; GE Healthcare). Pseudovirus entry routes were determined by pre-treating VeroE6-TMPRSS2 and Calu-3 cell monolayers with a concentration range of camostat mesylate or E64D diluted in Opti-MEM I (1×) + GlutaMAX (Gibco) for 2 hours prior to infection with 1 × 10^3^ pseudoviruses. The cells were incubated at 37°C overnight and scanned on the Amersham Typhoon Biomolecular Imager (channel Cy2; resolution 10 mm; GE Healthcare). All pseudovirus experiments were quantified using ImageQuant TL 8.2 image analysis software (GE Healthcare).

#### GFP complementation cell-cell fusion assays

The GFP complementation cell-cell fusion assay with transient spike protein expression was performed as previously described (*42*). Briefly, 1.5 μg pGAGGS-spike DNA and pGAGGS-β-Actin-P2A-7xGFP11-BFP DNA or empty vector DNA were transfected into HEK-293T cells with PEI (Polysciences) in a ratio of 1:3 (DNA: PEI). Transfected HEK-293T cells were transferred in three technical replicates to GFP1-10 expressing VeroE6, VeroE6-TMPRSS2 and Calu-3 cell monolayers in a ratio of 1:80 (HEK-293T cells : GFP1-10 expressing cells). Cell-cell fusion events were quantified at 18 hours post transfer using Amersham Typhoon Biomolecular Imager (channel Cy2; resolution 10 μm; GE Healthcare). Data was analyzed using the ImageQuant TL 8.2 image analysis software (GE Healthcare) by calculating the sum of all GFP+ pixels per well.

For the authentic virus GFP complementation cell-cell fusion assay Calu-3 cells expressing GFP1-10 and Calu-3 cells expressing GFP11 were seeded in a 1:1 ratio in 96-well Cell Carrier Ultra plates (Perkin Elmer) in and were incubated at 37°C in a humidified CO_2_ incubator until confluent. Virus was diluted in 2-fold serial dilutions starting with 20.000 viral particles based on 8-hour infectious viral titers in 200μl Opti-MEM I (1×) + GlutaMAX. 100μl of each dilution was added to the monolayers of 1 × 10^5^ Calu-3 cells. Cells were incubated with inoculums at 37°C for 24 hours in a humidified CO_2_ incubator and cells were fixed in 4% paraformaldehyde for 20 minutes. The cells were washed three times in PBS and stained for Hoechst 33343 for 30 minutes and washed again three times in PBS. Next, cells were imaged using the Opera phenix spinning disk confocal HCS system (Perkin Elmer) equipped with a 10x air objective (NA 0.3) and 405 nm and 488 nm solid state lasers. Hoechst and GFP were detected using 435-480 nm and 500-550 nm emission filters, respectively. Nine fields per well were imaged covering approximately 50% of the individual wells. The sum of GFP positive syncytia per well were quantified using the Harmony software (version 4.9, Perkin Elmer). Cellular origin of the GFP positive areas was confirmed by eye, using the Hoechst labeled nuclei.

#### Western blotting

SDS-PAGE and western blotting was performed as described before (*42*). Briefly, concentrated pseudovirus stocks were diluted to a final concentration of 1x Laemmli loading buffer (Bio-Rad) containing 5% 2-mercaptoethanol. Authentic viruses were diluted to a final concentration of 2x Laemmli loading buffer containing 5% 2-mercaptoethanol. All samples were boiled for 30 min at 95°C. Samples were used for SDS-PAGE analysis using precast 10% TGX gels (Bio-Rad). Gels and subsequent transfer onto 0.45 μm Immobilon-FL PVDF membranes in TGS containing 20% methanol. Spike was stained using mouse-anti-SARS-CoV-2 S2 (1:1000, Genetex), SARS-CoV-2 nucleoprotein was stained using rabbit-anti-SARS-CoV NP (1:1000, Sino Biological) and VSV nucleoprotein was stained using mouse-anti-VSV-N (1:1000, Absolute Antibody). All blots were subsequently stained by infrared-labelled secondary antibodies (1:20,000; Licor). Western blots were scanned on an Odyssey CLx and analyzed using Image Studio Lite Ver 5.2 software.

#### Statistical analysis

Statistical analysis was performed with the GraphPad Prism 9 software. Groups were compared by one- or two-way ANOVA, followed by a Benjamini and Hochberg (original FDR method; Q=0.05) or a Sidak multiple-comparison test, respectively, or by an unpaired T-test, as indicated in the figure legends.

**Fig. S1.**
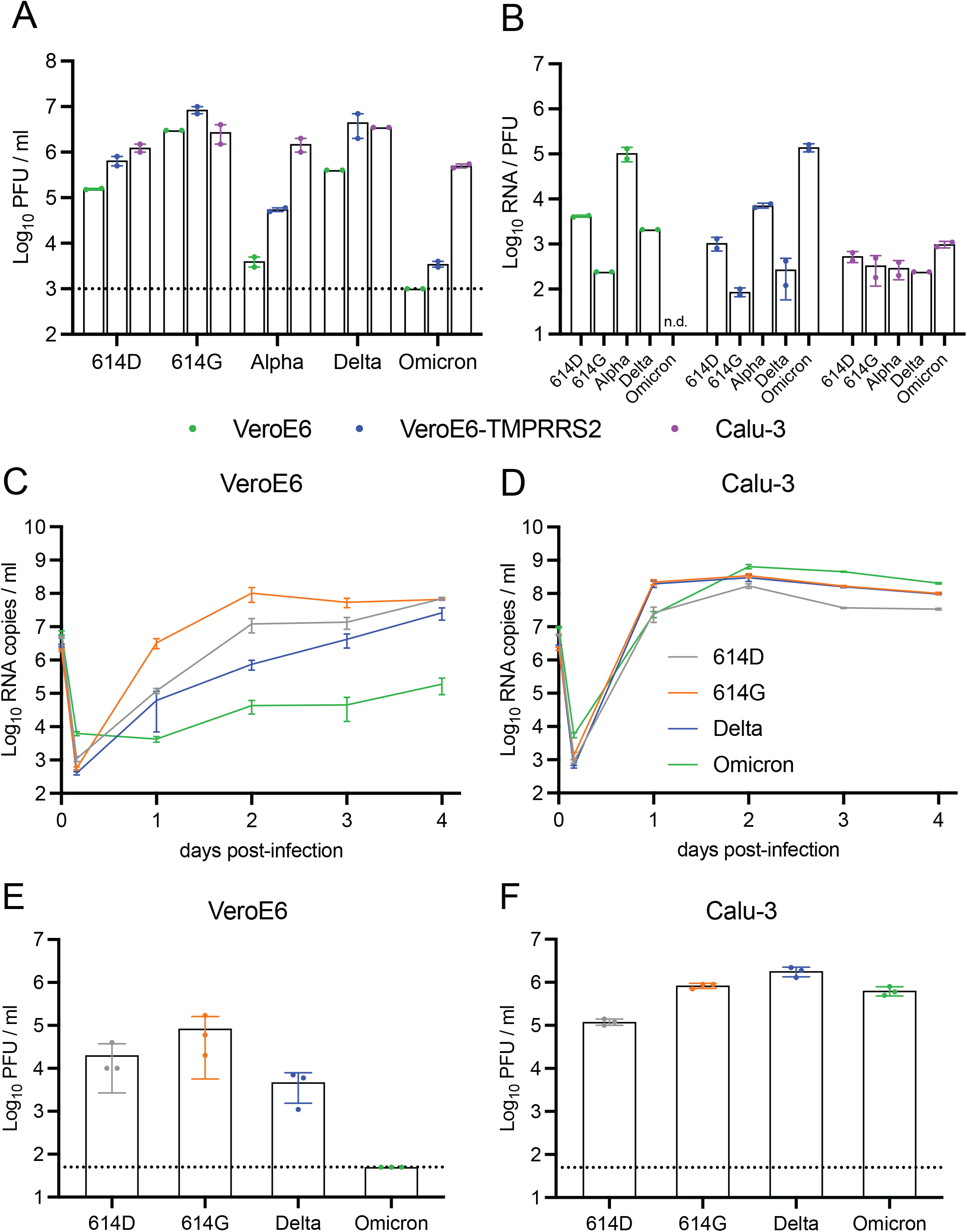
Omicron is attenuated on VeroE6 and VeroE6-TMPRSS2 cells. (a) Infectious virus titers of the 614D, 614G, Alpha, Delta and Omicron virus stocks were determined by calculating PFU on VeroE6, VeroE6-TMPRSS2 cells and Calu-3 cells. Dotted line indicates limit of detection. (b) The infectivity of 614D, 614G, Alpha, Delta and Omicron virus stocks was determined by calculation of the RNA copies to PFU ratio on VeroE6, VeroE6-TMPRSS2 and Calu-3 cells. (c-f) Replication kinetics of 614D, 614G, Alpha, Delta and Omicron in terms of RNA (c-d) and PFU at 2 d.p.i. (e-f) on VeroE6 cells (c and e) and Calu-3 cells (d and f). Dotted line indicates limit of detection. Error bars indicate SEM. PFU = plaque forming units.

**Fig. S2.**
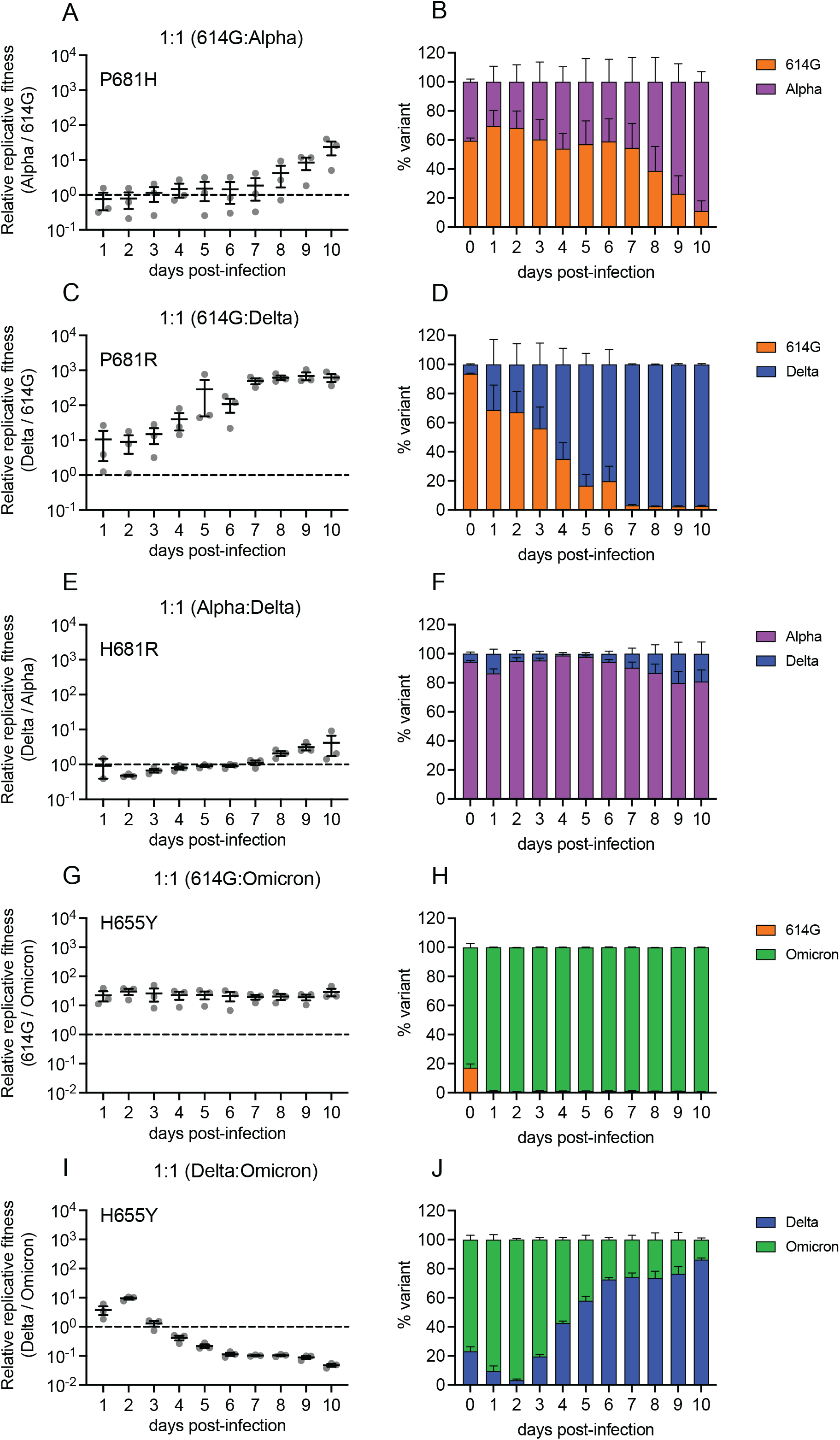
At a 1:1 ratio, Omicron has a fitness advantage over Delta in the first days and rapidly outcompetes 614G. (a-b) Relative replicative fitness (a) and proportion of each variant (b) in 2D airway-organoids infected with a 1:1 ratio of an ancestral 614G virus to Alpha. (c-d) Relative replicative fitness (c) and proportion of each variant (d) in 2D airway-organoids infected with a 1:1 ratio of an ancestral 614G virus to Delta. (e-f) Relative replicative fitness (e) and proportion of each variant (f) in 2D airway-organoids infected with a 1:1 ratio of Alpha to Delta. (g-h) Relative replicative fitness (g) and proportion of each variant (h) in 2D airway-organoids infected with a 1:1 ratio 614G to Omicron. (i-j) Relative replicative fitness (i) and proportion of each variant (j) in 2D airway-organoids infected with a 1:1 ratio of Delta to Omicron. Error bars indicate SEM.

**Fig. S3.**
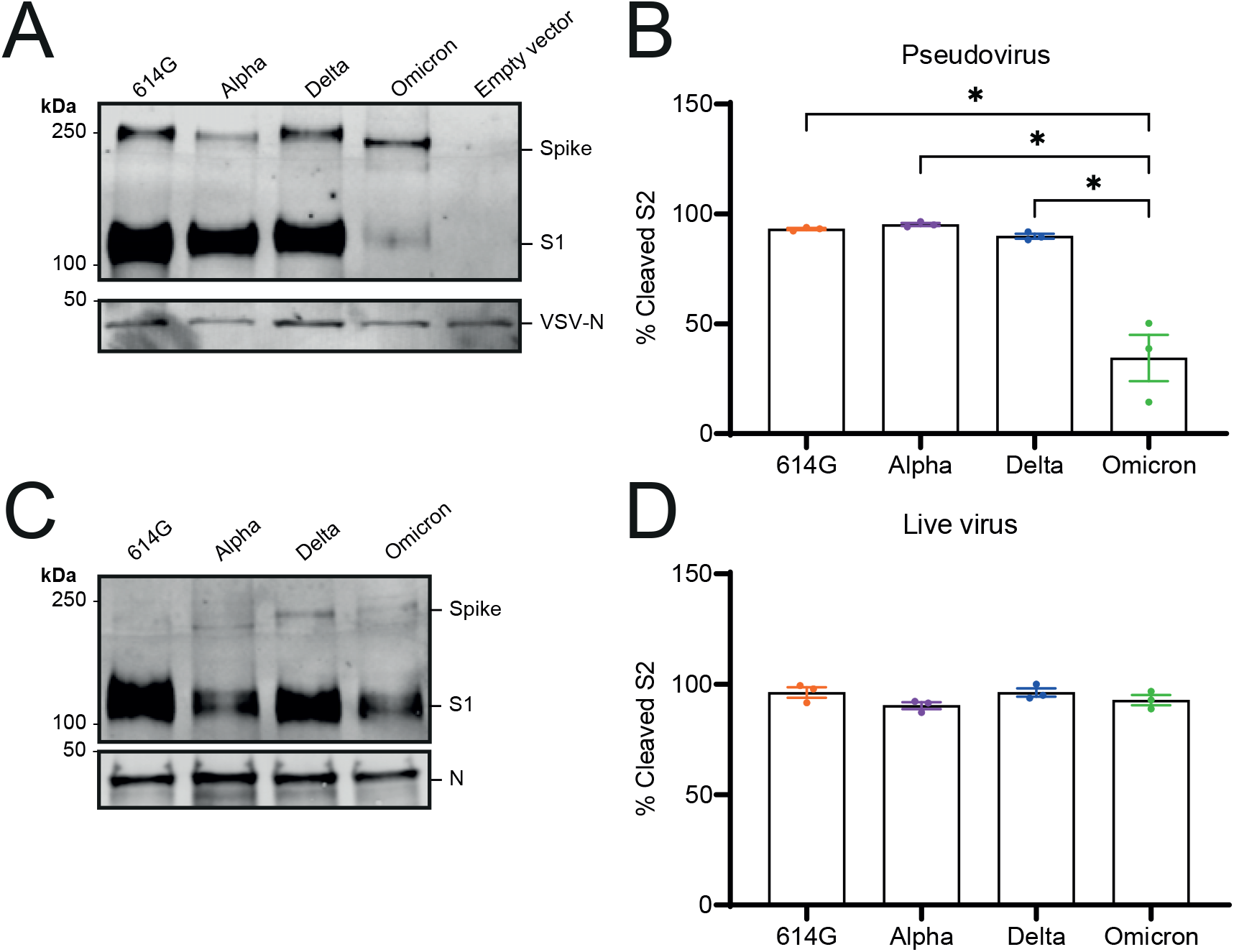
S1/S2 cleavage of pseudovirus and authentic virus stocks. (a-b) Immunoblotting of concentrated 614G, Alpha, Delta and Omicron pseudovirus stocks (a) and quantification (b). (c-d) Immunoblotting of 614G, Alpha, Delta and Omicron authentic virus stocks (c) and quantification (d). Groups were compared by one-way ANOVA. Error bars indicate SEM. * p < 0.05.

**Fig. S4.**
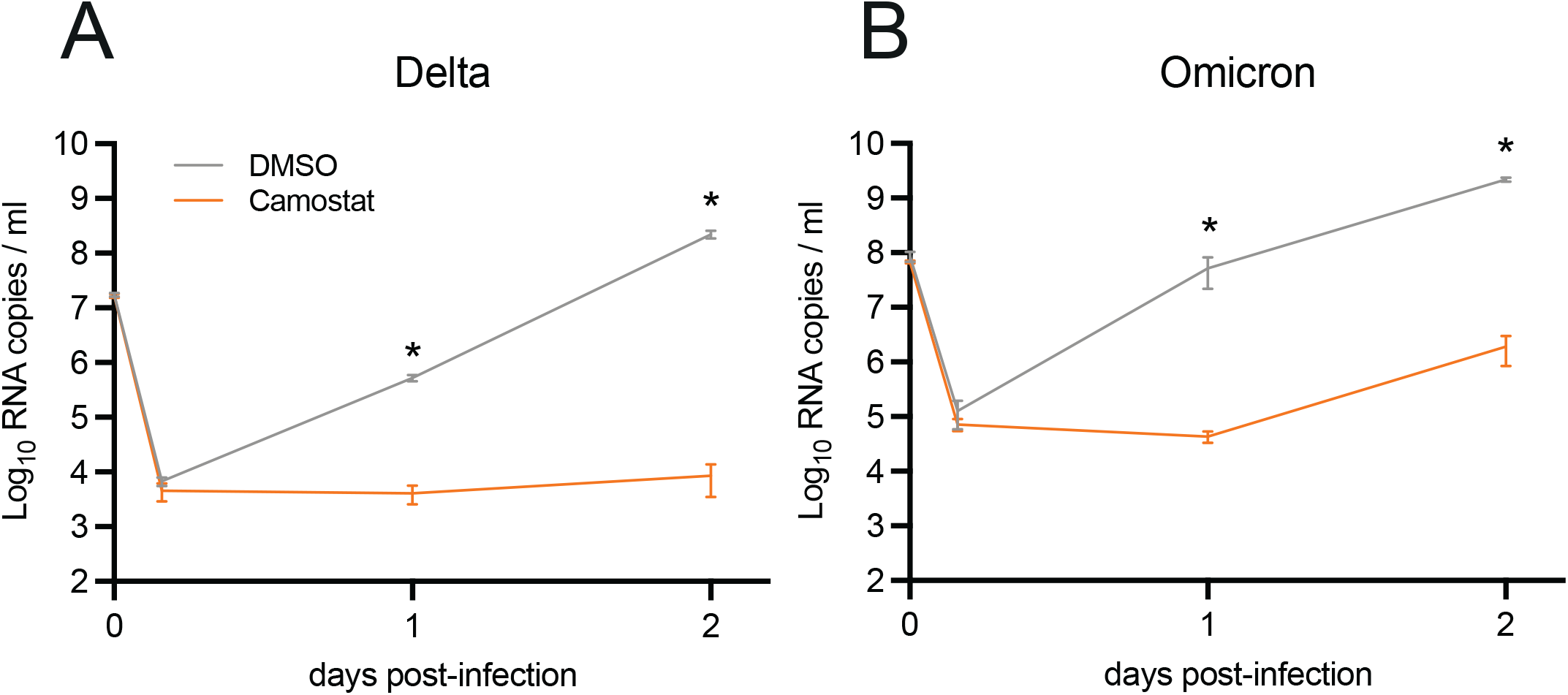
Delta and Omicron are inhibted by serine protease inhibitor camostat in air-liquid interface airway organoids. (a-b) Replication of Delta (a) and Omicron (b) in the presence of DMSO (vehicle) or camostat (100 μM). Groups were compared by two-way ANOVA. *p < 0.05. Error bars depict SEM.

**Fig. S5.**
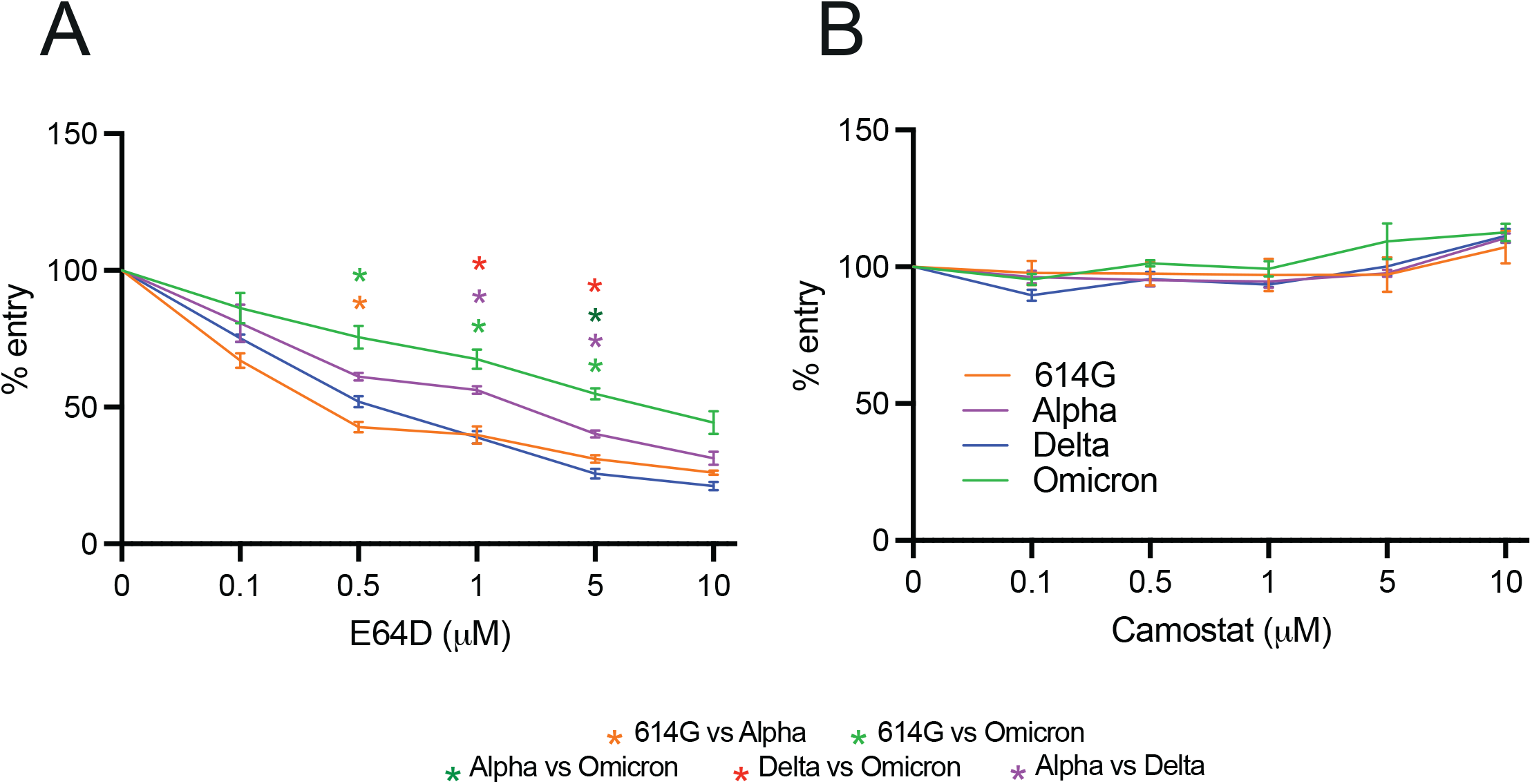
Entry into VeroE6 cells can be inhibited by E64D. (a-b) Percentage entry of pseudoviruses in VeroE6 cells pretreated with a concentration range of either E64D (a) or camostat (b). Groups were compared by two-way ANOVA. *p < 0.05. Error bars depict SEM.

**Fig. S6.**
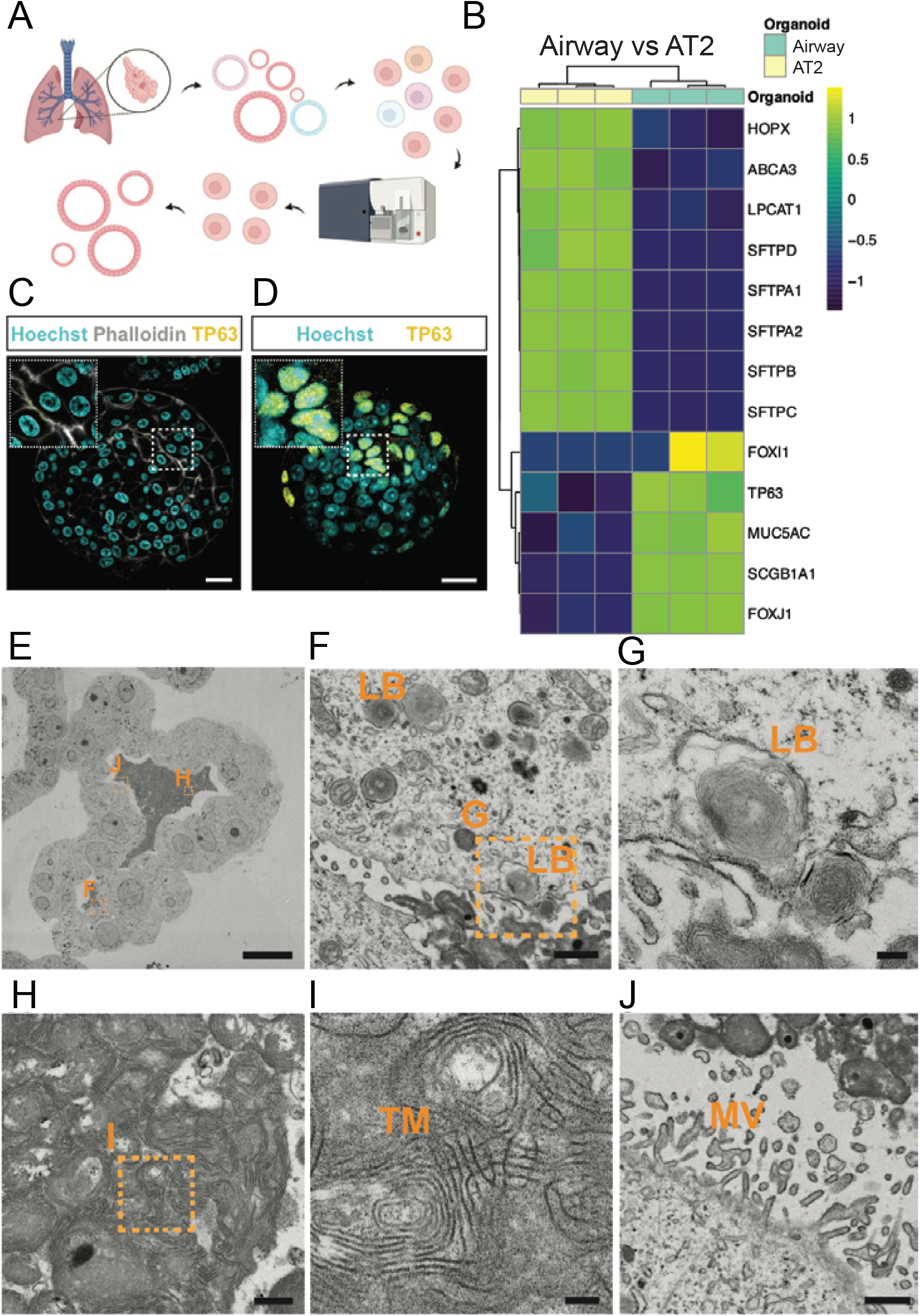
Generation and characterization of human adult alveolar type 2 (AT2) organoids. (a) Human adult stem cells are isolated from healthy lung tissue after lung resection surgery by enzymatic digestion of the tissue. Isolated stem cells are grown for 1 passage in basement membrane extract before they are dissociated and sorted for HTII-280 and Lysotracker high positivity to isolate a pure population of AT2 cells, which are again grown in basement membrane extract to generate AT2 organoids. (b) Heatmap of normalized expression values scaled across samples of selected airway and alveolar marker genes in airway and AT2 organoids. (c-d) Immunofluorescent staining of AT2 organoids (c) and airway organoids (d) for the basal cell marker TP63 (yellow). Nuclei are stained with hoechst (cyan) and actin is stained with phalloidin (white). (e-j) Overview of an intact alveolar type 2 (AT2) organoid (e) showing cytoplasmic lamellar bodies (LB) (f-g) and secreted lamellar bodies (h-i) that organize into tubular myelin (TM). (j) Apical microvilli (MV) are predominant on the AT2 cells. Scale bars indicate 20 μm (c-e), 1 μm (f-j), 200 nm (g), 500 nm (h), 100 nm (i). Panel (a) was created with BioRender.com.

**Fig. S7.**
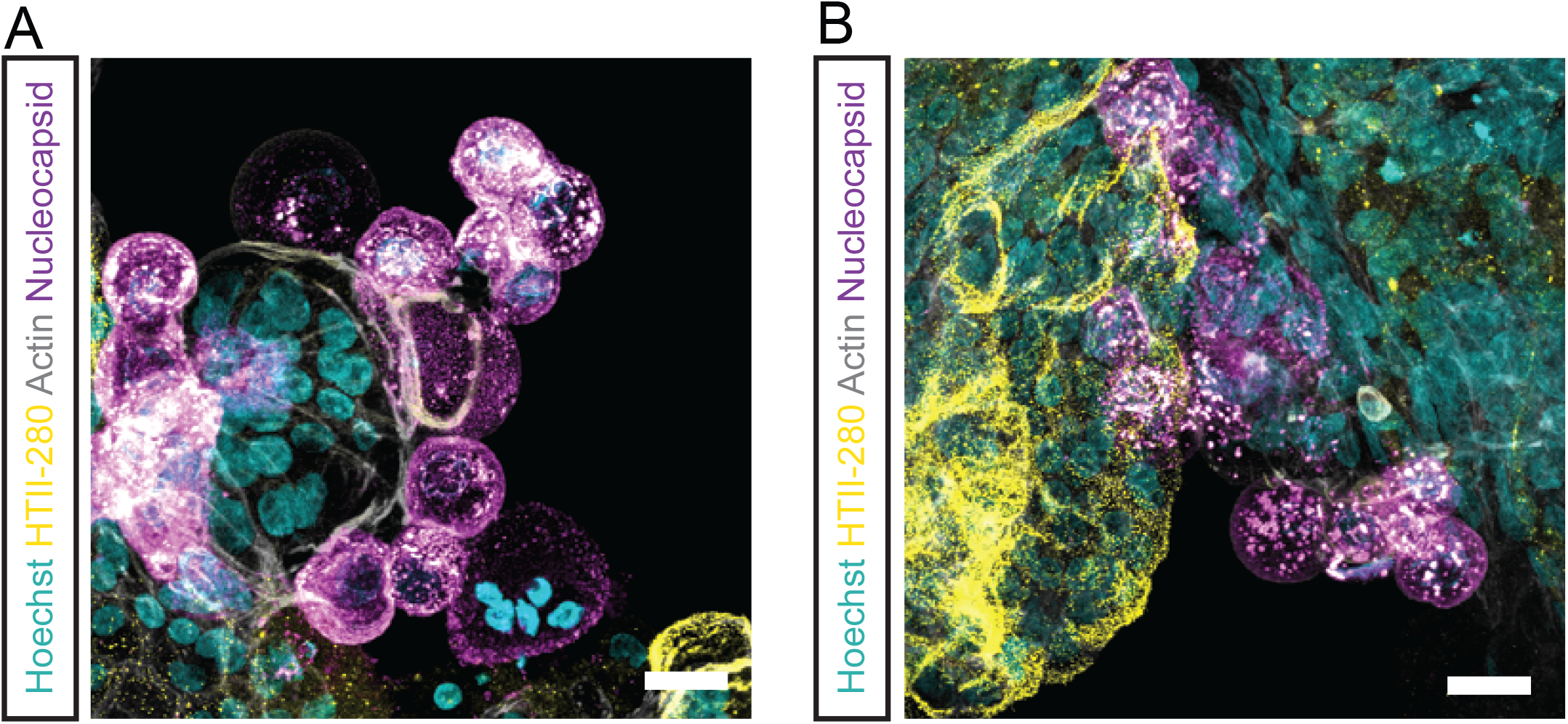
Confocal imaging of AT2 organoids at day 10 post-infection. (a-b) Immunofluorescent images of Delta (a) and Omicron (b) infected 3D AT2 cells at 4 d.p.i. Scale bars in a-b represent 20 μm.

## References

1. J. A. Plante et al., The variant gambit: COVID-19’s next move. Cell Host Microbe 29, 508–515 (2021).

2. K. Leung, M. H. Shum, G. M. Leung, T. T. Lam, J. T. Wu, Early transmissibility assessment of the N501Y mutant strains of SARS-CoV-2 in the United Kingdom, October to November 2020. Euro Surveill 26, (2021).

3. M. S. Graham et al., Changes in symptomatology, reinfection, and transmissibility associated with the SARS-CoV-2 variant B.1.1.7: an ecological study. Lancet Public Health, (2021).

4. D. Frampton et al., Genomic characteristics and clinical effect of the emergent SARS-CoV-2 B.1.1.7 lineage in London, UK: a whole-genome sequencing and hospital-based cohort study. Lancet Infect Dis, (2021).

5. E. Volz et al., Assessing transmissibility of SARS-CoV-2 lineage B.1.1.7 in England. Nature, (2021).

6. F. Campbell et al., Increased transmissibility and global spread of SARS-CoV-2 variants of concern as at June 2021. Euro Surveill 26, (2021).

7. H. Allen et al., Household transmission of COVID-19 cases associated with SARS-CoV-2 delta variant (B.1.617.2): national case-control study. Lancet Reg Health Eur 12, 100252 (2022).

8. J. M. Carreno et al., Evidence for retained spike-binding and neutralizing activity against emerging SARS-CoV-2 variants in serum of COVID-19 mRNA vaccine recipients. EBioMedicine 73, 103626 (2021).

9. H. Gu et al., Probable Transmission of SARS-CoV-2 Omicron Variant in Quarantine Hotel, Hong Kong, China, November 2021. Emerg Infect Dis 28, (2021).

10. in CDC COVID-19 Science Briefs. (Atlanta (GA), 2020).

11. S. Cele et al., SARS-CoV-2 Omicron has extensive but incomplete escape of Pfizer BNT162b2 elicited neutralization and requires ACE2 for infection. medRxiv, (2021).

12. E. Petersen et al., Emergence of new SARS-CoV-2 Variant of Concern Omicron (B.1.1.529) -highlights Africa’s research capabilities, but exposes major knowledge gaps, inequities of vaccine distribution, inadequacies in global COVID-19 response and control efforts. Int J Infect Dis 114, 268–272 (2022).

13. F. Graham, Daily briefing: Omicron coronavirus variant puts scientists on alert. Nature, (2021).

14. K. Ito, C. Piantham, H. Nishiura, Relative Instantaneous Reproduction Number of Omicron SARS-CoV-2 variant with respect to the Delta variant in Denmark. J Med Virol, (2021).

15. Nishiura H. et al., Relative Reproduction Number of SARS-CoV-2 Omicron (B.1.1.529) Compared with Delta Variant in South Africa. Journal of clinincal medicine 11, (2022).

16. CDC, COVID data tracker CDC. (2022).

17. W. F. Garcia-Beltran et al., mRNA-based COVID-19 vaccine boosters induce neutralizing immunity against SARS-CoV-2 Omicron variant. medRxiv, (2021).

18. A. M. Syed et al., Omicron mutations enhance infectivity and reduce antibody neutralization of SARS-CoV-2 virus-like particles. medRxiv, (2022).

19. Y. Wang et al., The significant immune escape of pseudotyped SARS-CoV-2 variant Omicron. Emerg Microbes Infect 11, 1–5 (2022).

20. L. Liu, e. al., Striking antibody evasion manifested by the Omicron variant of SARS-CoV-2. Nature, (2021).

21. J. M. Carreño, e. al., Activity of convalescent and vaccine serum against SARS-CoV-2 Omicron. (2022).

22. W. Dejnirattisai et al., Omicron-B.1.1.529 leads to widespread escape from neutralizing antibody responses. bioRxiv, (2021).

23. W. Dejnirattisai et al., Reduced neutralisation of SARS-CoV-2 omicron B.1.1.529 variant by post-immunisation serum. Lancet, (2021).

24. W. F. Garcia-Beltran et al., mRNA-based COVID-19 vaccine boosters induce neutralizing immunity against SARS-CoV-2 Omicron variant. Cell, (2022).

25. C. Geurts van Kessel, et al., Divergent SARS CoV-2 Omicron-specific T- and B-cell responses in COVID-19 vaccine recipients. MedRxiv, (2021).

26. M. M. Lamers et al., SARS-CoV-2 productively infects human gut enterocytes. Science 369, 50–54 (2020).

27. M. M. Lamers et al., Human airway cells prevent SARS-CoV-2 multibasic cleavage site cell culture adaptation. Elife 10, (2021).

28. M. M. Lamers et al., An organoid-derived bronchioalveolar model for SARS-CoV-2 infection of human alveolar type II-like cells. EMBO J 40, e105912 (2021).

29. J. J. A. van Kampen et al., Duration and key determinants of infectious virus shedding in hospitalized patients with coronavirus disease-2019 (COVID-19). Nat Commun 12, 267 (2021).

30. J. H. Ahn et al., Nasal ciliated cells are primary targets for SARS-CoV-2 replication in the early stage of COVID-19. J Clin Invest 131, (2021).

31. K. P. Y. Hui et al., Tropism, replication competence, and innate immune responses of the coronavirus SARS-CoV-2 in human respiratory tract and conjunctiva: an analysis in ex-vivo and in-vitro cultures. Lancet Respir Med, (2020).

32. J. A. Plante et al., Spike mutation D614G alters SARS-CoV-2 fitness. Nature 592, 116–121 (2021).

33. A. W. Byrne et al., Inferred duration of infectious period of SARS-CoV-2: rapid scoping review and analysis of available evidence for asymptomatic and symptomatic COVID-19 cases. BMJ Open 10, e039856 (2020).

34. X. He et al., Temporal dynamics in viral shedding and transmissibility of COVID-19. Nat Med 26, 672–675 (2020).

35. K. R. W. Emary et al., Efficacy of ChAdOx1 nCoV-19 (AZD1222) vaccine against SARS-CoV-2 variant of concern 202012/01 (B.1.1.7): an exploratory analysis of a randomised controlled trial. Lancet 397, 1351–1362 (2021).

36. A. S. V. Shah et al., Effect of Vaccination on Transmission of SARS-CoV-2. N Engl J Med 385, 1718–1720 (2021).

37. E. Callaway, Omicron likely to weaken COVID vaccine protection. Nature 600, 367–368 (2021).

38. C. Zeng et al., SARS-CoV-2 spreads through cell-to-cell transmission. Proceedings of the National Academy of Sciences 119, e2111400119 (2022).

39. J. Buchrieser et al., Syncytia formation by SARS-CoV-2-infected cells. EMBO J 40, e107405 (2021).

40. L. Jackson, e. al., SARS-CoV-2 cell-to-cell spread occurs rapidly and is insensitive to antibody neutralization. BioRxiv, (2022).

41. Q. Sattentau, Avoiding the void: cell-to-cell spread of human viruses. Nat Rev Microbiol 6, 815–826 (2008).

42. A. Z. Mykytyn et al., SARS-CoV-2 entry into human airway organoids is serine protease-mediated and facilitated by the multibasic cleavage site. Elife 10, (2021).

43. R. J. Hulswit, C. A. de Haan, B. J. Bosch, Coronavirus Spike Protein and Tropism Changes. Adv Virus Res 96, 29–57 (2016).

44. J. K. Millet, G. R. Whittaker, Host cell proteases: Critical determinants of coronavirus tropism and pathogenesis. Virus Res 202, 120–134 (2015).

45. M. Hoffmann et al., SARS-CoV-2 Cell Entry Depends on ACE2 and TMPRSS2 and Is Blocked by a Clinically Proven Protease Inhibitor. Cell 181, 271–280 e278 (2020).

46. J. Beumer, Geurts, M.H., Lamers, M.M., Puschhof, J., Zhang, J., van der Vaart, J., Mykytyn, A.Z., Breugem, T.I., Riesebosch, S., Schipper, D., van den Doel, P.B., de Lau, W., Pleguezuelos-Manzano, C., Busslinger, G., Haagmans, B.L., Clevers H., A CRISPR/Cas9 genetically engineered organoid biobank reveals essential host factors for coronaviruses. Nat Commun 12, (2021).

47. S. Matsuyama et al., Enhanced isolation of SARS-CoV-2 by TMPRSS2-expressing cells. Proc Natl Acad Sci U S A 117, 7001–7003 (2020).

48. J. Wei et al., Genome-wide CRISPR screen reveals host genes that regulate SARS-CoV-2 infection. bioRxiv, (2020).

49. J. E. Park et al., Proteolytic processing of Middle East respiratory syndrome coronavirus spikes expands virus tropism. Proc Natl Acad Sci U S A 113, 12262–12267 (2016).

50. C. Tarnow et al., TMPRSS2 is a host factor that is essential for pneumotropism and pathogenicity of H7N9 influenza A virus in mice. J Virol 88, 4744–4751 (2014).

51. N. Iwata-Yoshikawa et al., TMPRSS2 Contributes to Virus Spread and Immunopathology in the Airways of Murine Models after Coronavirus Infection. Journal of Virology 93, (2019).

52. T. Ebisudani et al., Direct derivation of human alveolospheres for SARS-CoV-2 infection modeling and drug screening. Cell Rep 35, 109218 (2021).

53. J. Youk et al., Three-Dimensional Human Alveolar Stem Cell Culture Models Reveal Infection Response to SARS-CoV-2. Cell Stem Cell, (2020).

54. A. A. Salahudeen et al., Progenitor identification and SARS-CoV-2 infection in human distal lung organoids. Nature, (2020).

55. H. Katsura et al., Human Lung Stem Cell-Based Alveolospheres Provide Insights into SARS-CoV-2-Mediated Interferon Responses and Pneumocyte Dysfunction. Cell Stem Cell, (2020).

56. C. Maslo et al., Characteristics and Outcomes of Hospitalized Patients in South Africa During the COVID-19 Omicron Wave Compared With Previous Waves. JAMA, (2021).

57. F. Abdullah et al., Decreased severity of disease during the first global omicron variant covid-19 outbreak in a large hospital in tshwane, south africa. Int J Infect Dis, (2021).

58. E. Mahase, Covid-19: Hospital admission 50-70% less likely with omicron than delta, but transmission a major concern. BMJ 375, n3151 (2021).

59. N. Wolter, et al., Early assessment of the clinical severity of the SARS-CoV-2 Omicron variant in South Africa. MedRxiv, (2021).

60. U. H. S. Agency, SARS-CoV-2 variants of concern and variants under investigation in England: technical briefing 33. Dec 2021. (2021).

61. D. Kim, J. Jo, J. S. lim, S. Ryu, Serial interval and basic reproduction number of SARS-CoV-2 Omicron variant in South Korea. MedRxiv, (2021).

62. W. S. Hart, e. al., Generation time of the Alpha and Delta SARS-CoV-2 variants. MedRxiv, (2021).

63. M. Diamond et al., The SARS-CoV-2 B.1.1.529 Omicron virus causes attenuated infection and disease in mice and hamsters. Res Sq, (2021).

64. N. Sachs et al., Long-term expanding human airway organoids for disease modeling. EMBO J 38, (2019).

65. T. Hashimshony et al., CEL-Seq2: sensitive highly-multiplexed single-cell RNA-Seq. Genome Biol 17, 77 (2016).

66. S. Simmini et al., Transformation of intestinal stem cells into gastric stem cells on loss of transcription factor Cdx2. Nat Commun 5, 5728 (2014).

67. H. Li, R. Durbin, Fast and accurate long-read alignment with Burrows-Wheeler transform. Bioinformatics 26, 589–595 (2010).

68. M. I. Love, W. Huber, S. Anders, Moderated estimation of fold change and dispersion for RNA-seq data with DESeq2. Genome Biol 15, 550 (2014).

69. Y. Benjamini, Y. Hochberg, Controlling the False Discovery Rate - a Practical and Powerful Approach to Multiple Testing. J R Stat Soc B 57, 289–300 (1995).

70. F. G. Faas et al., Virtual nanoscopy: generation of ultra-large high resolution electron microscopy maps. J Cell Biol 198, 457–469 (2012).

